# Patient-Derived Organoids as a Model to Understand Tumor Microenvironment-Driven Nano-Bio Interactions – A Framework Towards Improved Nanomedicine Translation

**DOI:** 10.64898/2026.09.03.749195

**Authors:** Meenu Priya Resmi, Jayati Chakrabarti, Kelvin W Pond, Swarna Ganesh

**Affiliations:** Department of Cellular and Molecular Medicine, University of Arizona College of Medicine, Tucson, Arizona, United States; Department of Biomedical Engineering, University of Arizona College of Engineering, Tucson, Arizona, United States; University of Arizona Cancer Center, Tucson, Arizona, United States; BIO5 Institute, University of Arizona, Tucson, Arizona, United States

## Abstract

Nanomedicines that perform well in conventional cell cultures often fail to translate into patients, in part due to the inability to reproduce the protein corona, and hence the biological identity, that nanoparticles acquire within the tumor microenvironment (TME). Here, we establish patient-derived colonic organoids (PDCOs) as a platform for profiling nano-bio interactions under physiologically tumor-relevant conditions using graphene quantum dots (GQD) and Silicon Quantum Dots (SiQD) as material-distinct probes. By combining spectral flow cytometry, confocal imaging, dynamic light scattering, and multi-spectral dimension-reduction analysis of single-cell uptake, we resolve how nanoparticle identity, uptake, and intracellular fate diverge between conventional culture and patient tissue. Conventional two-dimensional culture overestimated internalization by two-fold relative to ex vivo PDCOs, while TME-specific proteins directed nanoparticles to distinct organoid subpopulations. Exploiting this, we engineer the protein corona with tumor-specific proteins to redirect SiQDs to chemoresistant cells, from 3.8% to 70%, an 18-fold increase in targeted delivery to the cells that drive therapy resistance. Internalization was also model-dependent. PDCOs and fibroblasts favored clathrin/dynamin (DNM1)-mediated endocytosis into lysosomes, whereas cancer cells favored caveolae (CAV1)-mediated entry that bypasses lysosomal degradation. This distinction determines whether a nanocarrier is degraded or delivered intact. Collectively, these findings turn the protein corona into a programmable design tool and establish PDCOs as a next-generation platform for precision nano-oncology.

**Graphical Abstract:** 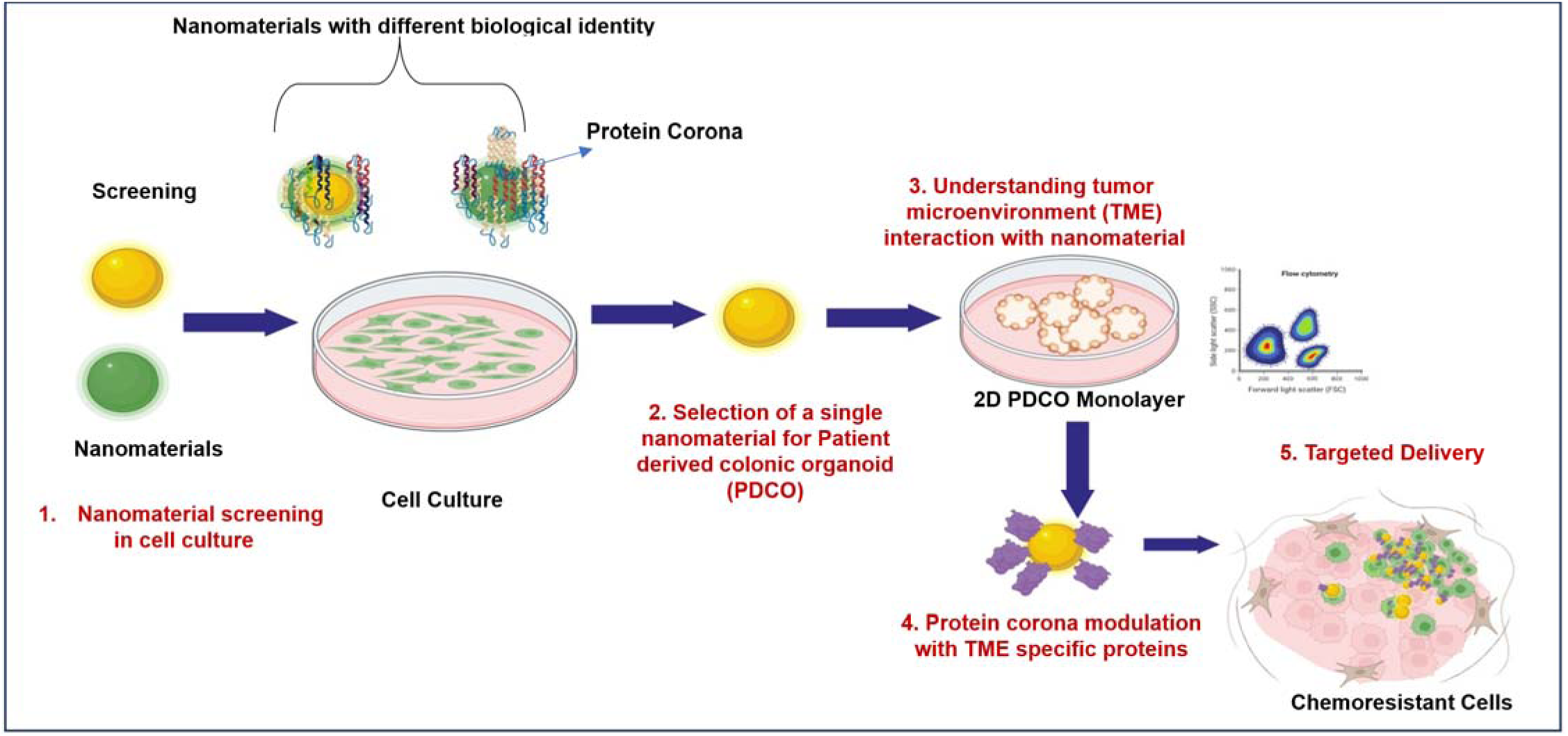

## Introduction

Nanoparticles offer a comprehensive strategy for tackling major challenges in cancer treatment, including targeted drug delivery ^1^, high-precision imaging ^2^, and therapeutics ^3^. Despite significant progress in nanomedicine research, only a limited number of nanoparticles that succeed in in vitro models advance to clinical trials, since they do not yield the same outcome in actual patients ^4^. The primary reason for this translational gap is the tumor’s heterogeneity, which affects how nanoparticles accumulate, localize, and traffic within cells. The intracellular fate of the nanoparticle determines tumor-targeting efficiency and off-target effects, which ultimately influence therapeutic outcome ^5^.

Due to the inability to recreate the interactions directly observed in patients, nanomedicine research has historically relied on two-dimensional cell culture, and animal models have been fundamental to deepening our understanding of nano-bio interactions and enabling nanoparticle screening ^6^. However, two-dimensional cell cultures are simplified and do not capture the full physiological complexity of tumors. While preclinical animal models can provide improved physiological conditions, cross-species variation in serum proteins, immunological responses, and tumor physiology remains a critical challenge. Research shows that responses to nanomedicine often vary between cell cultures, animal models, and clinical settings ^7^, showing that nanoparticles assume unique identities depending on the biological environment.

Research indicates that the differences in biological identities observed in both in vitro and in vivo models stem from their distinct protein-corona formation ^8, 9^. When a nanoparticle is introduced into a biological environment, proteins adsorb onto its surface to form a protein corona. This corona masks the nanoparticle surface and confers on a new biological identity that substantially influences the selective uptake and intracellular trafficking of nanoparticles, thereby impacting their clinical effectiveness ^10^.

The protein corona presents both an obstacle and an opportunity. To limit the effects of the protein corona, studies have developed nanoparticles from core-crosslinked block copolymer micelles, but these have reduced long-term stability ^11^ and have been coated with a dense hydrophilic shell to prevent protein interactions ^11^. However, such coatings can potentially induce immunogenicity and may impair targeting capabilities ^12^. Moreover, eliminating the corona is not always desirable, as it can also promote nanoparticle transport and cellular uptake ^13^.

Rather than merely preventing the protein corona, current research focuses on understanding and fine-tuning it to program the nanoparticle for cellular uptake ^14^. Most research so far has focused on characterizing the serum-derived protein corona, which includes abundant proteins such as apolipoproteins ^15^, immunoglobulins, albumin ^16^, and complement proteins ^17^. However, there is limited research on how tumor microenvironment (TME) proteins affect biological identity and cell interactions ^18^. These knowledge gaps hinder precise predictions of nano-bio interactions in patients, potentially affecting clinical outcomes.

Efforts to address this gap have utilized three-dimensional spheroids ^19^ and organoids ^20^ to investigate nano-bio interactions within the TME. These studies provided important insights into nanoparticle delivery, therapeutic responses, and mechanisms of action using models that better mimic physiological conditions. However, these three-dimensional models were derived from established cell lines and did not fully capture patient-specific heterogeneity. Consequently, the impact of patient-derived proteins on the biological identity that affects nanoparticle applications remains largely underexplored.

We therefore hypothesized that this gap in understanding reflects a specific, testable premise: that the protein corona formed in serum-based culture differs from that formed under patient-relevant TME conditions, and that these differences drive measurably different nanoparticle uptake and intracellular trafficking. Patient-derived organoids provide a complementary model between immortalized cell lines and animal studies by preserving donor-derived epithelial features within an experimentally tractable culture system. Human colorectal organoids can be expanded from normal, adenomatous, and malignant tissue while retaining important genetic and phenotypic characteristics of the tissue of origin. In selected gastrointestinal cancers, patient-derived organoids have also reproduced clinically observed differences in treatment response, supporting their use as translational preclinical models while recognizing that epithelial-only organoids do not reproduce stromal, immune, or vascular components of the complete tumor microenvironment ^21^. We previously showed that primary human colonic organoids cultured as two-dimensional monolayers self-organize into crypt-like epithelial tissues containing spatially distinct stem-like, proliferative, and differentiated compartments ^22^ ^23^ ^24^. This organization, combined with uniform access to nanoparticles and compatibility with high-content imaging and flow cytometry, provides a controlled system for determining how epithelial heterogeneity influences nano-bio interactions.

Here we introduce a framework that integrates the PDCOs for screening nanomedicines with the vision of clinical translation. Using graphene quantum dots (GQD) and silicon quantum dots (SiQD) as representative nanoparticles, we performed multimodal quantitative profiling of nanomaterial uptake across in vitro models (fibroblasts and cancer cells) and ex vivo PDCOs to probe the nano-bio interface and systematically evaluate TME-specific interactions. To keep all models comparable in dimensionality, we used a 2D monolayer of patient-derived colorectal polyp organoids that retains key features of native colonic tissue, with distinct stem-like, proliferative, and differentiated populations organized into crypt-like node and non-node compartments, providing a physiologically relevant platform for studying nanoparticle uptake in human colonic epithelium^21^.

Using this framework, we show that the biological identity of nanoparticles, and hence the uptake and intracellular fate, is governed by both material composition and the surrounding biological environment. SiQD and GQD form distinct protein coronas and are internalized differently across fibroblasts, cancer cells, and PDCOs, with traditional cell culture overestimating internalization compared with ex vivo models. Further, engineering the protein corona with tumor microenvironment-specific proteins (Wnt and EGF) selectively directs SiQD to chemoresistant cell populations (CD44v9), showing that protein corona composition can be used to target therapy–resistant populations. Pathway analysis further revealed that PDCOs and conventional in vitro models favor different endocytic pathways for nanoparticle internalization, with corresponding differences in intracellular trafficking. Together, these findings confirm that tumor microenvironment-specific proteins govern nanoparticle uptake and targeting and position PDCOs as a next-generation platform for precision nano-oncology, aiming to support personalized nanomedicine that maximizes therapeutic targeting while minimizing off-target effects.

## Results and Discussion

### Characterization of the SiQD and GQD Protein Corona Interaction

Previous research has shown that the protein corona can significantly influence nanoparticle cellular internalization ^8, 25^. Studies have shown that the protein corona formation depends on the nanoparticle physicochemical characteristics, including size, shape, charge, and surface chemistry ^26, 27^. We used SiQD and GQD as model systems to test the hypothesis. The size (<20nm) and spherical shape were maintained to assess material-specific properties in protein corona formation. Additionally, we strategically selected physicochemical properties to maximize protein adsorption in biological environments, given their high surface-to-volume ratio ^28, 29^. To confirm the unique surface chemistry of the quantum dots, Fourier Transform Infrared Spectroscopy (FT-IR) was performed, and it indicated that SiQD has Si-N, Si-H, and Si-O-Si bonds on its surface ^30^. Meanwhile, GQD shows C=O and CH_2_ stretches ^31^, reflecting the primary differences in surface chemistry **(Supplementary Figure 1)**. To investigate and confirm protein adsorption, we incubated QDs in complete growth medium for 90 minutes. When a QD is placed in a biological environment, higher-affinity proteins interact with it, forming a tightly bound layer around the nanoparticle ^32^ known as the hard protein corona. Soft corona proteins are low-affinity proteins that also bind to the QD surface but are highly unstable. To assess the higher-affinity proteins, a hard protein corona was isolated by centrifuging the QD protein complex at high speed for 45 min, and the supernatant containing lower-affinity proteins was removed and quantified separately ^33^ **(Figure 1A)**. To confirm the protein corona formation and understand the protein corona thickness, the size distribution of SiQD and GQD protein corona was compared using transmission electron microscopy **(Supplementary Figure 2)**.

**Figure 1:**
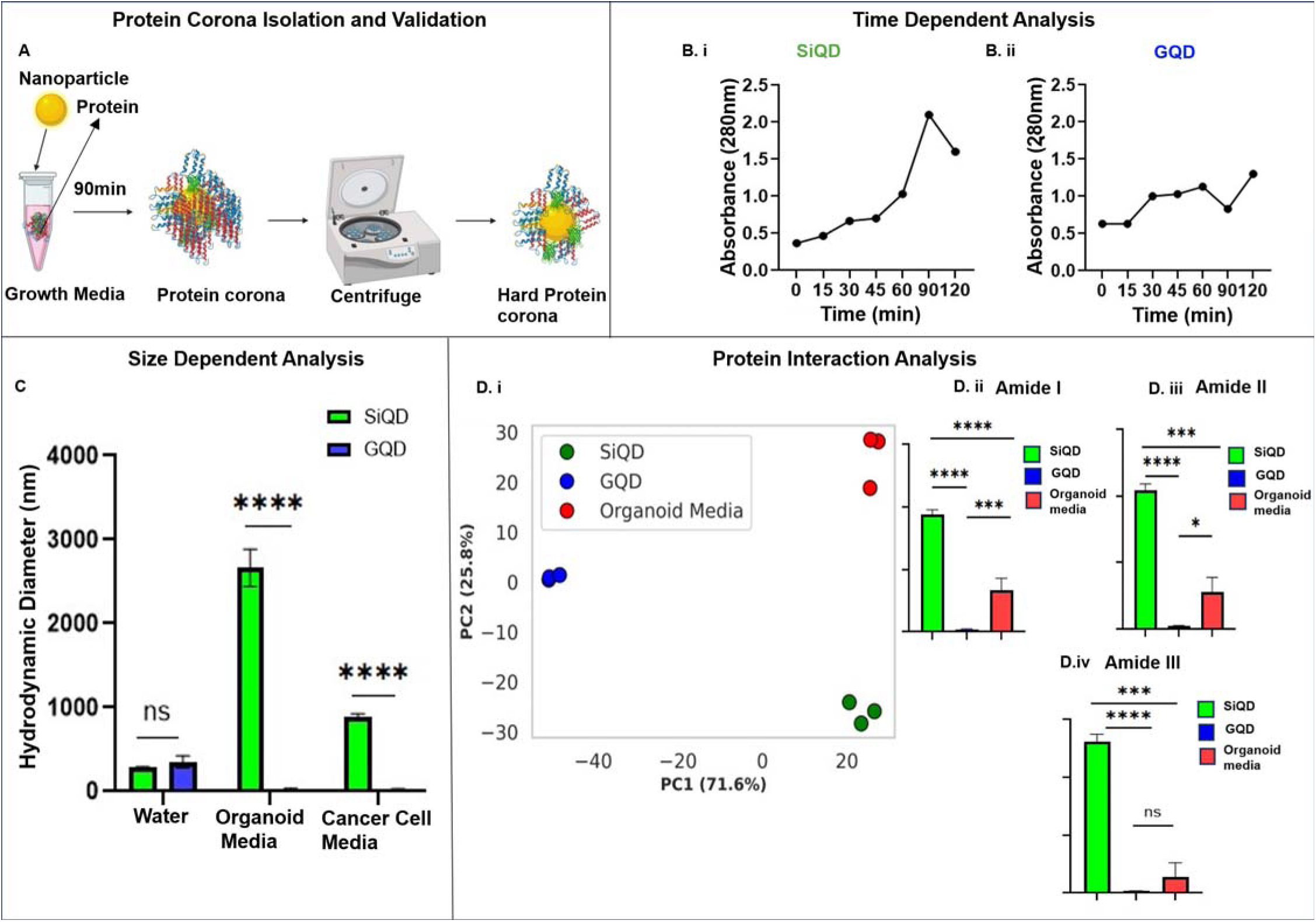
**A.** Representation shows the Isolation of the protein corona from growth media. **B. i, ii** shows the concentration of the protein corona formation by the SiQD and GQD in Fetal Bovine Serum at 0,15,30,45,60,90 and 120 minutes. **C.** Comparison of the hydrodynamic diameter of SiQD and GQD in water, Organoid Media, and Cancer Cell Media using DLS, n = 3, t-test. **D. i** Principal component analysis comparing the protein structure in organoid growth media, SiQD, and GQD protein corona formed i organoid growth media using FTIR, n=3, t-test. **D. ii**, **iii, iv** Bar plots representing the comparison of Amide I, II, III between SiQD, GQD, and organoid Media, n=3, t-test. Statistical significance: P < 0.05*, P < 0.01**, P < 0.001***, P < 0.00001****, P>0.05 **ns**

To mechanistically study protein interactions on SiQDs and GQDs, we adopted a three-phase approach. In the first phase, we used fetal bovine serum (FBS), a major source of proteins and growth factors in most traditional cell culture, to investigate the material-dependent dynamics of protein corona formation. To investigate this, SiQDs and GQDs were incubated in FBS at 15-minute intervals for 2 hours. In the case of SiQDs, the maximum corona formed within 90 minutes, after which protein desorption from the SiQDs was observed **(Figure 1B. i)**. Similarly, GQDs showed peak protein adsorption within 60 minutes, followed by desorption at 90 min; after that, an increase in protein adsorption was observed that could be attributed to the aggregation of the QDs **(Figure 1B. ii)**. Adsorption and desorption of the protein by the QDs were consistent with the Vroman effect driving the process ^34^. When a QD interacts with the biological environment, highly abundant proteins bind rapidly at first, and later higher-affinity proteins displace them until equilibrium is reached ^35^. These observations confirm that protein corona formation is dynamic and differs significantly between SiQDs and GQDs.

The next phase is to incorporate more physiologically relevant conditions, including cancer cell media and organoid media. Cancer cell medium contains a basal medium with amino acids, glucose, vitamins, and salts, supplemented with 10% FBS, which approximates simplified physiological conditions. The organoid media contain growth factors, signaling cues, and proteins that support the growth of heterogeneous cell populations and better mimic physiological conditions ^36, 37^ **(Supplementary Table 1)**. To account for these diverse biological environments, we incubated the QDs in cancer cell medium and organoid medium. Protein corona formation determines the colloidal stability of the QDs. Depending on the material, the protein corona can either cause aggregation or prevent it. We used dynamic light scattering (DLS) to characterize QD behavior across different biological environments by measuring hydrodynamic diameter. SiQD and GQD showed markedly different behavior in biological environments. In water, both GQD and SiQD are aggregated. After incubation with cancer cell and organoid media, SiQD’s hydrodynamic diameter significantly increased **(Supplementary Figure 3)**. When a protein interacts with a QD, the strong interaction can induce structural changes in the protein, revealing additional binding sites. This allows further interactions with other proteins, leading to aggregation ^38^.

Conversely, the hydrodynamic diameter of GQD was smaller compared to its size in water **(Supplementary Figure 4)**, mainly because the protein corona caused steric repulsion ^39^, which improved the colloidal stability of GQD. Furthermore, we compared the hydrodynamic diameters of SiQD and GQD in the growth media. In Water, SiQD and GQD have hydrodynamic diameters of 288.6 nm and 353.2 nm, respectively, indicating that GQD and SiQD are aggregated in water. In organoid media, SiQD measures over 2665 nm, whereas GQD is only 27.9 nm. In cancer cell media, SiQD is 891.5 nm, and GQD is 79.28 nm. These results suggest that SiQD tends to aggregate due to the formation of a protein corona, while protein corona formation protects GQD from aggregation **(Figure 1C)**. This confirms that protein adsorption patterns and aggregation behavior differ significantly between GQD and SiQD.

FT-IR was used to study protein backbone interactions, enabling comparison of protein interactions at the molecular level with SiQD and GQD **(Supplementary Figure 5)**. We analyzed the protein backbone, including Amide I, Amide II, and Amide III bonds at 1650 cm-1, 1550 cm-1, and 1400 cm-1, respectively. In SiQDs, significant differences in the Amide I, II, and III bands relative to organoid media are observed, indicating a change in the overall protein structure upon interaction **(Figure 1D. ii)**. By contrast, GQDs also showed significant differences in Amide I and II but not in Amide III **(Figure 1D. iii)**, confirming that GQD and SiQD interact differently with proteins in the biological environment. To evaluate the consistency of protein corona interactions and assess variations in the structural interactions of QDs with proteins, Principal component analysis was performed on multiparametric FT-IR data.

Principal component analysis (PCA) of FT-IR spectra has shown that PC1 and PC2 contribute 71.6% and 25.8% of the variance, respectively. The PCA score plot showed that GQD, SiQD, and organoid media tightly clustered along each axis **(Figure 1D. i)**. Overall, PCA shows that GQD and SiQD have no overlap, and protein corona formation is consistent across samples. The protein adsorption patterns, aggregation behavior, and structural interactions with proteins indicate that the overall composition of the material determines protein corona formation. This confirms that GQD and SiQD will exhibit distinct biological identities due to protein corona formation, which contributes to variable uptake in vitro and in vivo.

### Profiling of GQD and SiQD Uptake in Normal and Cancerous Cell Lines

Figure 1 shows that GQD and SiQD exhibit variable biological identities due to protein corona formation. The next step is to determine whether these variable biological identities drive cellular interactions. To investigate this, we quantified uptake, profiled phenotypes, assessed uptake heterogeneity, and evaluated toxicity. We used a fibroblast (LRFB) and a colon cancer cell line (COLO201). QDs exhibit fluorescence from quantum confinement, which we tracked via flow cytometry and confocal microscopy. SiQDs and GQDs emit fluorescence at 530nm and 421nm, respectively. We observed increased fluorescence intensity after incubating COLO201 and LRFB with SiQDs and GQDs. This confirms SiQD and GQD uptake in both cell populations (Figure 2Ai**, ii, iii, iv)**.

**Figure 2:**
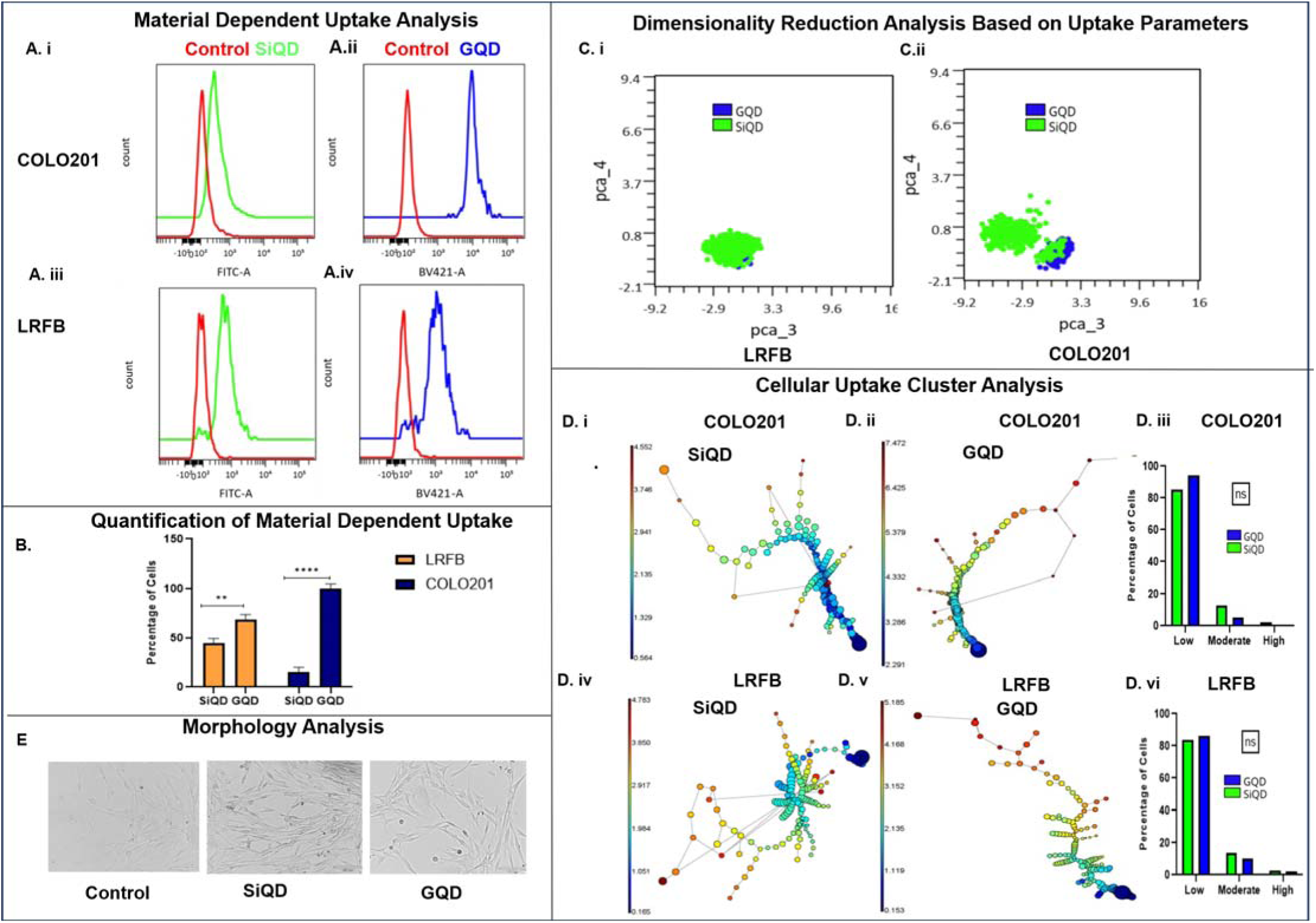
**A. i, ii, iii, iv**. Histogram representing the uptake of SiQD and GQD in COLO201 and LRFB. **B.** Bar plots showing the comparison of SiQD and GQD in COLO201 and LRFB based on the percentage of cell uptake, n= 10,000 cells. t-test**. C. i, ii.** PCA analysis analyzing the variabilities in SiQD and GQD positive cell populations in COLO201 and LRFB. **D. i, ii.** Cluster analysis of SiQD and GQD in COLO201. **D. iii.** Comparison of the percentage of cell populations of identified clusters ranked based on the intensity of SiQD and GQD uptake in COLO201. **D. iv, v.** Cluster analysis of SiQD and GQD in LRFB. **D. v.** Comparison of the percentage of cell populations of identified clusters ranked based on the intensity of SiQD and GQD uptake in LRFB.**E. i.** Brightfield image shows the morphology of the LRFB after incubating with SiQD and GQD, scale bar = 250um.Statistical significance: P < 0.05*, P < 0.01**, P < 0.001***, P < 0.00001****, P>0.05 **ns**

To understand the efficiency of SiQD and GQD uptake in each cell type, fluorescent intensity was quantified. In LRFB and COLO201 cells, GQD shows significantly higher fluorescence than SiQD, suggesting predominant accumulation of GQD within the cell population **(Supplementary Figure 6)**. Further, population analysis was performed to compare the percentage of QD-positive cells. In LRFB, GQD (68.66%) and SiQD (44.21%) were observed. Whereas in COLO201, GQD (99.91%) and SiQD (15.23%) were observed. This implies that GQDs exhibited higher uptake in colon cancer cell lines and fibroblasts (**Figure 2B**). These findings confirm that SiQD uptake is cell-specific, whereas GQD uptake is not cell-type-dependent, indicating a more generalized uptake pattern.

To compare material-specific uptake attributable to SiQD and GQD accumulation, we performed principal component analysis on the multidimensional flow cytometry dataset. PCA revealed that PC3 and PC4 were attributable to QD fluorescence intensity, which drives uptake **(Supplementary Figures 7 and 8)**. Further, the distribution of QDs in each cell line was compared along with these principal components. In fibroblasts, no significant variability in uptake was observed among the QDs, showing comparable internalization **(Figure 2C. i)**; in contrast, in colon cancer cell lines, a clear separation was observed between the SiQD and GQD **(Figure 2C. ii)**. These findings confirm that SiQD and GQD have material-specific uptake in cancer cells, whereas fibroblasts are less susceptible to material-specific QD uptake.

Further, to investigate heterogeneity in SiQD and GQD uptake in colon cancer cell lines and fibroblasts, we performed unsupervised clustering using fluorescence intensity and forward-and side-scattering, which reflect uptake, cell size, and complexity, respectively. The branching patterns ^40^ of SiQDs and GQDs in normal and cancer cell lines differ **(Figure 2. Di, ii, iv, v)**. This indicates differences in spatial clustering, confirming heterogeneity in QD uptake across samples. We further categorized these clusters as low-, moderate-, or high-fluorescence-intensity QD populations in each cell line. Comparing the distribution of fluorescence intensity, LRFB and COLO201 showed no variability in uptake **(Figure 2D. iii, vi).**

Furthermore, morphology-based analysis revealed no significant differences in morphology after incubation with SiQD or GQD compared to the control **(Figure 2E. i)**. This confirms that GQD and SiQD are biocompatible.

Altogether, this study compares SiQD and GQD internalization in both fibroblasts and cancer cells. Figure 2 shows that SiQD and GQD exhibited different uptake patterns in cancer cell lines. SiQD showed more variable, cell-type-specific uptake, whereas GQD showed more consistent uptake. We chose SiQD for subsequent experiments because its variable uptake suggests greater potential to accumulate in different cell populations, and this property can be further exploited for targeted cancer drug delivery, personalized therapeutics, improved imaging, and theranostic applications.

### Internalization of SiQD in Patient-Derived Colonic Organoids

The preceding experiments showed that SiQD and GQD association differs between fibroblasts and immortalized colorectal cancer cells, indicating that nanoparticle uptake depends on both material composition and cellular context. To further understand the tropism of specific nanoparticles for distinct cell types we evaluated SiQD behavior in an established 2D patient-derived colonic organoid (PDCO) model that preserves greater cellular and spatial heterogeneity and self-organizes into crypt-like node and non-node regions containing distinct stem-like, proliferative, and differentiated populations^22, 23,41, 24^. The two-dimensional configuration provides uniform nanoparticle exposure and enables direct confocal imaging, highly quantitative single-cell analysis, and cell recovery for flow cytometry while retaining epithelial lineage organization across tens of thousands of single cells.

To evaluate SiQD association with this patient-derived epithelium, organoid monolayers were exposed to SiQD for 24 hours and analyzed by bright-field microscopy, confocal z-stack imaging, and spectral flow cytometry **(Figure 3A)**. Sequential optical sections and orthogonal projections were used to determine whether SiQD fluorescence was present within the cellular volume rather than restricted to the monolayer surface. Flow cytometry then quantified SiQD-associated cells and the distribution of fluorescence across the heterogeneous organoid population. Together, these complementary measurements characterized SiQD association within a spatially organized human epithelial model and provided a basis for subsequent analysis of uptake heterogeneity among organoid subpopulations.

**Figure 3:**
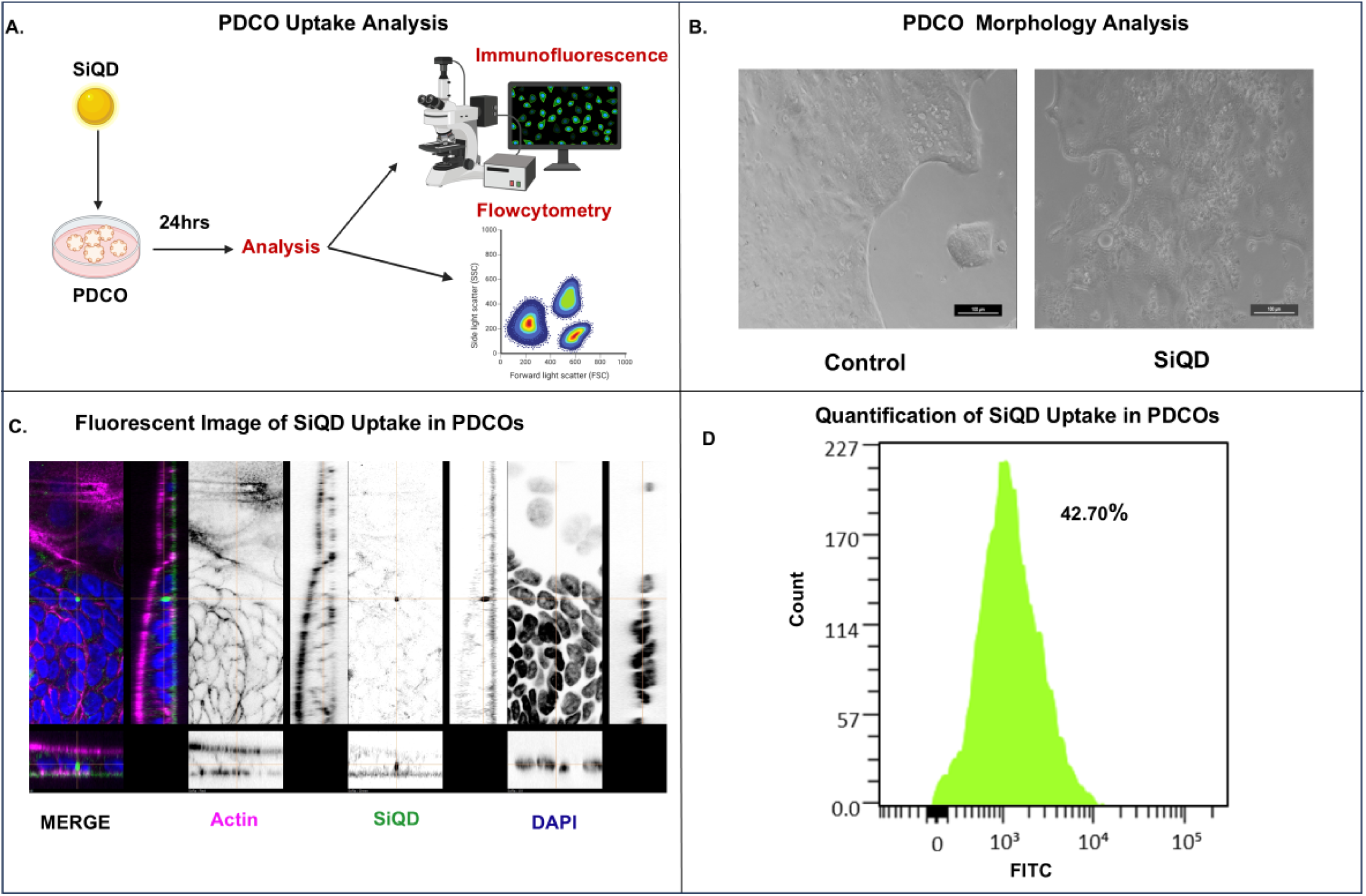
**A**. Schematic diagram of the SiQD uptake study in patient-derived colonic organoids (PDCO). **B**. Bright-field image of organoid control and SiQD-incubated organoids, scale bar = 100µm. **C**. Immunofluorescence analysis showing the uptake of SiQD in PDCO using Dapi, SiQD, Actin. **D. i**. Histogram represents the fluorescent intensity of SiQD between incubated and control PDCOs, Red = control, Green = SiQD incubated PDCO, n= 10,000 cells. **D. ii.** Histogram shows the SiQD-positive cell populations in PDCOs, n= 10,000 cells.

To confirm PDCO uptake, we performed immunofluorescence staining on SiQD-incubated patient-derived organoids **(Figure 3C).** Confocal microscopy revealed SiQD accumulation in PDCOs. To determine whether SiQD localizes intercellularly or on the cell surface, we performed sequential Z-stack imaging. We observed SiQD fluorescence at each focal plane, confirming effective uptake by the cells. This indicates that SiQD localizes within cells rather than on the organoid’s periphery.

To quantify SiQD internalization in PDCOs, we used spectral flow cytometry. A notable rise in fluorescence was observed in SiQD-incubated PDCOs, confirming the uptake **(Supplementary Figures 9 and 10)**. Population analysis showed 42.70% SiQD uptake in PDCOs **(Figure 3D)**. This study confirms that the SiQD uptake is relatively high in PDCOs.

To assess the structural integrity of the organoids after SiQD incubation, we evaluated morphology using bright-field microscopy. The bright-field image of the PDCO after 24 hr of SiQD incubation showed no significant difference in organoid morphology **(Figure 3B).**

Overall, conventional 2D cell culture does not capture the complexity of the tumor microenvironment. Even in vivo studies are limited in clinical translation due to divergent gene regulatory and signaling networks, spatial organization, and cell–cell interactions^42^. Using patient-derived organoids to understand nanoparticle biointeractions would better predict nanomedicine outcomes in clinical settings. Therefore, PDCOs can serve as a platform for customizing nanoparticles to enhance precision and translational potential.

### Temporal and Phenotypic Characterization of SiQD Internalization in Patient-Derived Colonic Organoids

In the previous figure, we confirmed and quantified SiQD uptake in PDCOs. Unlike cell cultures, the cell populations in PDCOs are highly heterogeneous, varying in granularity, size, and structural organization ^43^. Therefore, to better understand how SiQDs accumulate in physiologically relevant models, we characterized SiQD uptake by analyzing their distribution within PDCOs, examining correlations between SiQD internalization and cellular physical properties, and evaluating SiQD uptake kinetics in PDCOs.

To understand SiQD uptake dynamics in PDCOs, we performed unsupervised clustering based on fluorescence intensity, cell size, and cell complexity. (**Figure 4A i**) shows the cluster tree, colored by SiQD fluorescence intensity, confirming variability among cell populations. This analysis identified three main clusters of cells with similar features in size, granularity, and SiQD accumulation **(Figure 4A. ii)**. We then analyzed the distribution of SiQD uptake within each cluster. Cluster 1 (89.10%) has the highest percentage of cells that are more closely related, based on size, granularity, and intensity, followed by clusters 2 (9.18%) and 3 (1.73%). (Figure **4A** **ii).** We measured SiQD accumulation, cell size, and granularity in each cluster to understand how cellular heterogeneity in PDCOs affects uptake. Notably, Cluster 1 shows the highest SiQD uptake, cell complexity, and cell size, while Clusters 2 and 3 have lower values **(Figure 4B. i**). These results indicate a positive correlation between SiQD uptake, cell complexity, and cell size within these clusters.

**Figure 4:**
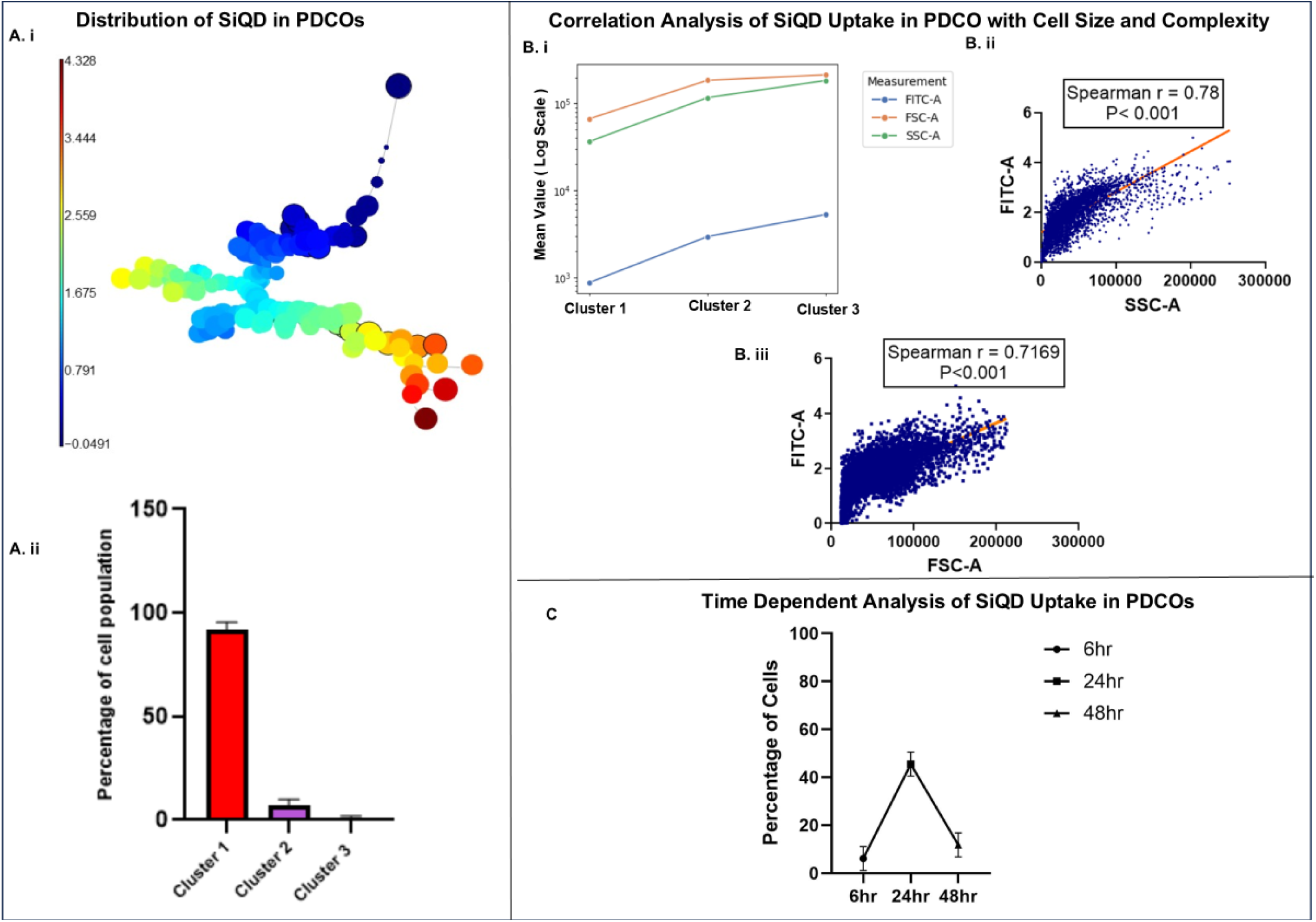
**A. i**. Clustering analysis based on the SiQD uptake, cell size, and complexity; each dot is colored based on the fluorescent intensity due to SiQD uptake. **A. ii** Bar plots show the percentage of each cell population in each cluster. **B. i**. Line plot connecting each cluster based on the cell size (FSC), cell complexity (SSC), and SiQD(FITC) intensity. **B. ii, B. iii**, **C.** Shows the percentage of SiQD uptake in organoids at 6, 24, and 48 hr; n = 10,000 cells.

Consequently, the study suggests that differences in cell size and granularity among organoid subpopulations contribute to variations in SiQD uptake affinity. Further research is needed to understand the underlying mechanism of specific SiQD uptake in PDCO subpopulations.

Previous research on traditional cell cultures has shown that QD uptake correlates directly with cell size and granularity ^44^. Cells with larger surface areas tend to have higher endocytic activity and greater uptake. To determine if this mechanism applies to PDCOs, the study examined the relationship between cell size, structural complexity, and fluorescence intensity using Spearman’s rank correlation. Specifically, we calculated the correlation coefficients between cell size (FSC), granularity (SSC), and SiQD fluorescence intensity. The coefficients for cell complexity versus uptake intensity and cellular size versus uptake intensity were 0.78 and 0.78169, respectively **(Figure 4B. ii, B. iii)**, indicating a monotonic relationship between cell size or complexity and SiQD uptake. This suggests that heterogeneity in cell size and granularity influences differential SiQD uptake among PDCO subpopulations.

To understand the temporal dynamics of SiQD internalization in PDCOs, we conducted a time-course analysis of SiQD uptake at 6, 24, and 48 hours. We measured the percentage of SiQD uptake at each time point. Uptake was evident at 6 hours, then increased sharply at 24 hours, and declined at 48 hours **(Figure 4C)**. This indicates that the highest SiQD uptake occurs at 24 hours, with a reduction possibly due to exocytosis at 48 hr ^45^. The study confirms that SiQD levels in this PDCO plateaued at 24 hours. Figure 4 shows that nanoparticle uptake depends on cell size, granularity, and time. Therefore, understanding how these factors affect uptake is crucial in PDCOs to optimize nanoparticles for better therapeutic responses, improved drug delivery efficiency, and higher predictability in clinical settings.

### Systemic Evaluation of Uptake Behavior of SiQD Across Various In Vitro Physiological Models

Figure 4 shows QD uptake in PDCOs. Next, we analyzed uptake to evaluate variations in nanobio interactions, from simplified traditional cell culture to complex PDCOs. We performed spectral flow cytometry on colon cancer cells, PDCOs, and fibroblasts after incubation with SiQD to examine differences in uptake patterns **(Figure 5A)**. We then compared uptake efficiency and kinetics.

**Figure 5:**
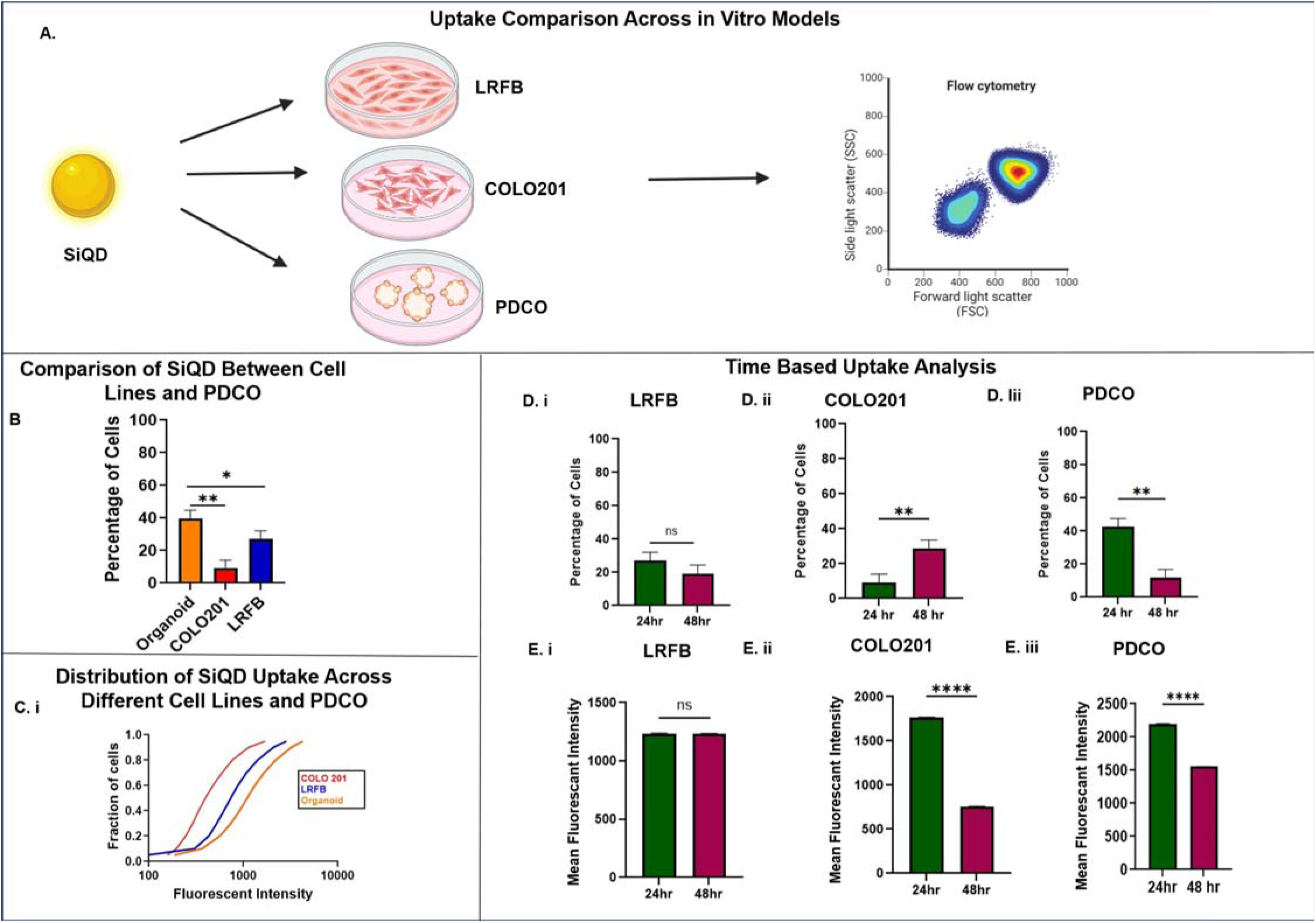
**A**. Schematic representation of the comparison of the SiQD uptake in LRFB, COLO201 and PDCO. **B**.Barplot compares the percentage of uptake across LRFB, COLO201 and PDCOs, n= 10,000 cells,t test. **Ci**, Empirical cumulative fraction of analysis of SiQD uptake in COLO201, LRFB and organoid. **D. i, ii, iii** Barplot representing the comparison of percentage of uptake at 24 and 48 hr in LRFB, COLO201 and PDCO, respectively, n= 10,000 cells, t-test. **E. i, ii, iii** Mean fluorescent intensity exhibited by SiQD at 24 hr and 48 hr in LRFB, COLO201 and PDCO, respectively, n= 10,000 cells, t-test. Statistical significance: P < 0.05, P < 0.01, P < 0.001, P < 0.00001

We assessed uptake efficiency for each cell type by measuring fluorescence intensity and the percentage of uptake after 24 hours of SiQD incubation. PDCOs showed a higher percentage of uptake compared to LRFB and COLO201 **(Figures 5B. i, ii)**. Additionally, the overall fluorescent intensity of the SiQD-posit ve cell population revealed that organoids accumulate more SiQD than LRFB and COLO201 **(Supplementary Figure 11).**

To assess how SiQD uptake is distributed across cell populations, we performed empirical cumulative functional analysis of FITC intensity at each percentile of cell populations in COLO201, LRFB, and patient-derived organoids **(Figure 5C. i)**. This non-parametric analysis determines the proportion of cell uptake by the SiQD. This analysis indicated that organoids display a pronounced increase relative to COLO201 and LRFB, confirming that organoids have a higher fluorescent threshold across cumulative cell fractions and thus higher internalization efficiency. This study confirms that SiQD uptake varies significantly among cancer cells, fibroblasts, and PDCOs.

To analyze uptake kinetics across fibroblasts, colon cancer cells, and PDCOs, we conducted time-dependent studies at 24 and 48 hours for each cell type. We quantified and compared uptake percentage and fluorescent intensity at each time point. In fibroblasts, there was no significant difference in uptake percentage and mean fluorescent intensity between 24 and 48 hours, **(Figures 5D. i, 5E. i)**, indicating no significant difference in intracellular uptake efficiency over time. In COLO201 cells, SiQD uptake increased, but intracellular accumulation significantly decreased at 48 hours **(Figures 5D. ii, 5E. ii)**. This suggests that while fewer SiQDs are retained at 48 hours, their distribution within cells becomes more dispersed due to a phenomenon called exocytosis. In PDCOs, both uptake percentage and fluorescent intensity increased at 24 hours and significantly decreased at 48 hours due to exocytosis **(Figures 5D. iii, 5E. iii)**. During exocytosis, cells actively transport nanoparticles from their interior to the outside environment 43. Overall uptake kinetics across cell types indicate that PDCOs, fibroblasts, and cancer cells exhibit significantly different kinetics.

This study shows that cancer cells and normal cells have significantly different uptake rates compared to PDCOs. This distinction helps explain why nanomedicines often fail in trials and underscores that nanoparticles’ unpredictable behavior in clinical practice stems from the lack of physiologically relevant models.

### Manipulating the Protein Corona with the TME Related Protein for Targeted Delivery of SiQD to Sub Cell Population in PDCOs

Previous findings in this study show that nanoparticle uptake efficiency and their intracellular fate depend on the tumor microenvironment. The next step is to investigate if TME-related proteins influence nanoparticle uptake in PDCOs. Research indicates that intracellular proteins tend to replace serum proteins in the protein corona due to the vroman effect, thereby aiding QD uptake ^46^. However, the role of external proteins in the growth medium remains unclear, as they serve as a substrate for other proteins to bind to when nanoparticles are introduced into cells or PDCO cultures. Current studies often overlook how proteins in the growth media influence protein corona formation, which can affect nanoparticle uptake in vitro models. Interactions. To address this, SiQDs were pre-engineered with cancer cell and organoid growth media, and their uptake was assessed to determine whether the growth media influence nano-bio interactions in PDCOs. In cancer cell growth media, the primary protein source is FBS, which is rich in serum proteins such as albumin, transferrin, immunoglobulins, and apolipoproteins. In contrast, organoid growth media mainly contain proteins derived from the tumor microenvironment **(Supplementary Table 1)**. Protein corona formation in conventional cell culture therefore differs substantially from that in patient-derived organoids, with the latter more closely reflecting the tumor microenvironment. Initially, we examined the behavior of SiQDs in both organoid growth media and cancer cell growth media (DMEM + FBS) through a size-and time-dependent analysis. To compare nanoparticle size in the growth media, DLS was performed 6 h after incubating SiQDs in cancer cell growth media and organoid growth media. The size of the SiQD protein corona in organoid media (4850 nm) was markedly larger than in cancer cell media (2384 nm), suggesting that SiQDs aggregate more in organoid media owing to protein crosslinking ^47^ **(Supplementary Figure 3)**. To further study protein corona formation dynamics, SiQDs were incubated in growth media, and adsorbed proteins were isolated at 15-minute intervals over two hours and their concentrations measured. In cancer cell growth media, the protein corona peaked at 65 minutes and then dissociated through the Vroman effect, whereas in organoid media maximum adsorption occurred at 75 minutes before decreasing **(Supplementary Figure 12)**. These findings demonstrate that SiQD behavior differs between the two media, consistent with the DLS results, and indicate that SiQD develops a distinct biological identity in the presence of TME-related proteins compared with serum proteins.

The distinct protein corona formed in organoid and cancer cell growth media prompted us to examine how this biological identity affects interactions with the tumor microenvironment. SiQD was incubated in cancer cell media and organoid media for 90 minutes and then added to PDCOs. Uptake was assessed using spectral flow cytometry, and SiQD uptake was quantified and categorized **(Figure 6A)**. The mean fluorescence intensity of SiQD-positive cells was measured, and SiQD preincubated in organoid media showed greater accumulation in organoids than SiQD preincubated in cancer cell media **(Figure 6B. i)**. However, the percentage of cell uptake did not differ significantly between the two conditions **(Figure 6B. ii)**. Together, these results indicate that differences in the protein corona substantially affected SiQD uptake in PDCOs.

**Figure 6.**
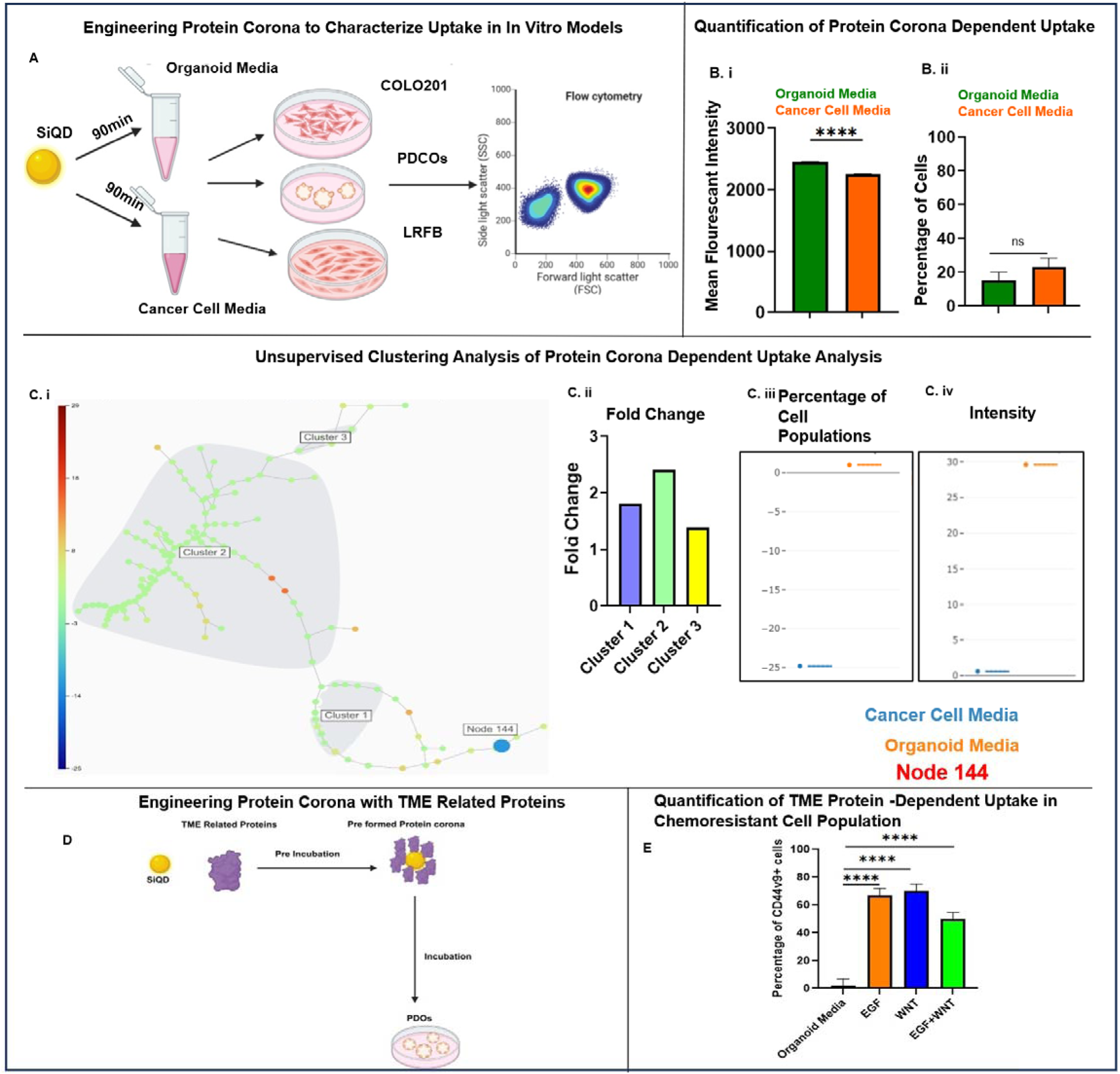
**A**. Schematic representation of the engineered protein corona to characterize uptake in cancer cells, fibroblasts, and PDOs. **B. i, ii**. Quantification of the SiQD uptake based on mean fluorescent intensity and percentage of cells, n=10,000 cells, t-test. **C. i**, Treeplot representing the uptake of SiQD after preincubating cancer cell media with organoid media, n= 10,000 cells. **C. ii**, Fold change of the SiQD fluorescent intensity across clusters between pre-organoid media and pre-cancer cell media.C. iii, iv, percentage of cell population and fluorescent intensity of SiQD preincubated with organoid media and cancer cell media in node 144, n= 10,000 cells. **D**. Schematic representing the engineering of the SiQD protein with TME-related media to study the targeted uptake of SiQD in PDOs. **E**. Percentage of chemoresistant cell uptake of SiQD after pre-engineering the protein corona with EGF, Wnt, and EGF + Wnt. N= 10,000 cells, t-test. Statistical significance: P < 0.05*, P < 0.01**, P < 0.001***,P < 0.00001****

These differences in uptake across conditions confirm the role of TME-related proteins. To examine them further, the pre-incubated organoid and cancer cell growth media samples were combined to analyze SiQD distribution among cell populations. Merging the datasets identified three common clusters based on SiQD accumulation, cell size, and granularity, where each node represents a cluster of cells **(Figure 6C. i)**. Fold changes in uptake intensity per cluster showed that pre-incubation in cancer cell media led to greater accumulation than pre-organoid conditions **(Figure 6C. ii)**. We then selected the node with the highest cell count that did not belong to clusters 1, 2, or 3: node 144, which differs markedly from the other clusters, contains over 30% of cells from pre-organoid conditions and shows higher SiQD fluorescence intensity (**Figure 6C. iii, iv)**. This indicates that cells within node 144 contribute substantially to overall intensity. Overall, organoid media promotes the specific accumulation of SiQD in certain cell types that cancer cell media do not, showing that TME-specific proteins in the organoid growth media regulate SiQD uptake in a specific subset of PDCO cell populations.

Having confirmed that TME-related proteins in the organoid growth media facilitate the selective accumulation of SiQD, we next asked whether deliberately manipulating the protein corona with a defined TME protein could target nanoparticles to a specific subset of the organoid cell population. From the organoid media, Wnt and epidermal growth factors (EGF) were selected for their established roles in chemoresistance. Wnt, a secreted glycoprotein, binds Frizzled and LRP5/6 receptors and activates the Wnt /β-catenin pathway, driving stem cell-like behavior and uncontrolled growth associated with resistance ^48^. EGF, a growth factor, binds to the Epidermal Growth Factor Receptor (EGFR) and activates RAS/RAF/MEK/ERK and PI3K/AKT pathways, promoting tumor growth, survival, and resistance ^49^. Wnt receptors and EGFR are highly expressed in CD44v9-positive chemoresistant cells ^50, 51^. To determine whether these TME proteins influence SiQD targeting, SiQDs were pre-incubated with Wnt, EGF, or Wnt + EGF, then washed by centrifugation to remove unbound protein before uptake was assessed in CD44v9-positive chemoresistant cells **(Figure 6D)**. Relative to complete media (3.8%), pre-incubation with Wnt (70%), EGF (66.85%), and Wnt + EGF (49.5%) produced a significant increase in uptake within the CD44v9-positive cell population **(Figure 6E)**. Because the engineered particles were washed before dosing, this increase reflects the particle-bound corona rather than free Wnt or EGF signaling.

Overall, this study shows that TME-related proteins in organoid growth media lead to selective accumulation of SiQDs in chemoresistant cell populations. Additionally, a Wnt-and EGF-based protein corona guides the nanoparticle to the CD44v9 cell population. This indicates the potential of using TME-related proteins for targeted tumor delivery, which could minimize off-target effects and improve biodistribution in cancer treatments.

### Comparison of SiQD Uptake Pathway Between Cancer Cells, Normal Cells, and PDCOs

Previous results have indicated variability in the uptake of different cell types. To explore whether the SiQD uptake pathway varies among cancer cells, normal cells, and PDCOs, a pathway analysis was performed, which is crucial for clinical translation. This study examines two main pathways that are dominant and well-studied for nanoparticle uptake by cells: clathrin-mediated and caveolae-mediated endocytosis. In clathrin-mediated endocytosis, clathrin and adaptor proteins form coated pits that enclose cargo molecules into vesicles. Dynamin (DNM1) then facilitates vesicle pinching from the membrane. These vesicles subsequently fuse with early endosomes, mature into late endosomes, and finally merge with lysosomes, leading to degradation^52^. In caveolin-mediated uptake, Caveolin 1 and 2 (CAV) promote caveolae formation, often aiding albumin accumulation. Once formed, caveolae pinch off and fuse with endosomes. Notably, the caveolin pathway can bypass lysosomes and direct vesicles to the endoplasmic reticulum ^53^**(Figure 7A)**. For nanoparticles, if they follow the clathrin pathway and reach lysosomes, they can be used for pH-res onsive drug delivery or targeted lysosomal cancer therapies. However, delivering proteins or genes to lysosomes may result in their degradation. Therefore, to deliver genes or proteins, nanoparticles must be fine-t ned to escape endosomes before lysosomal fusion.

**Figure 7:**
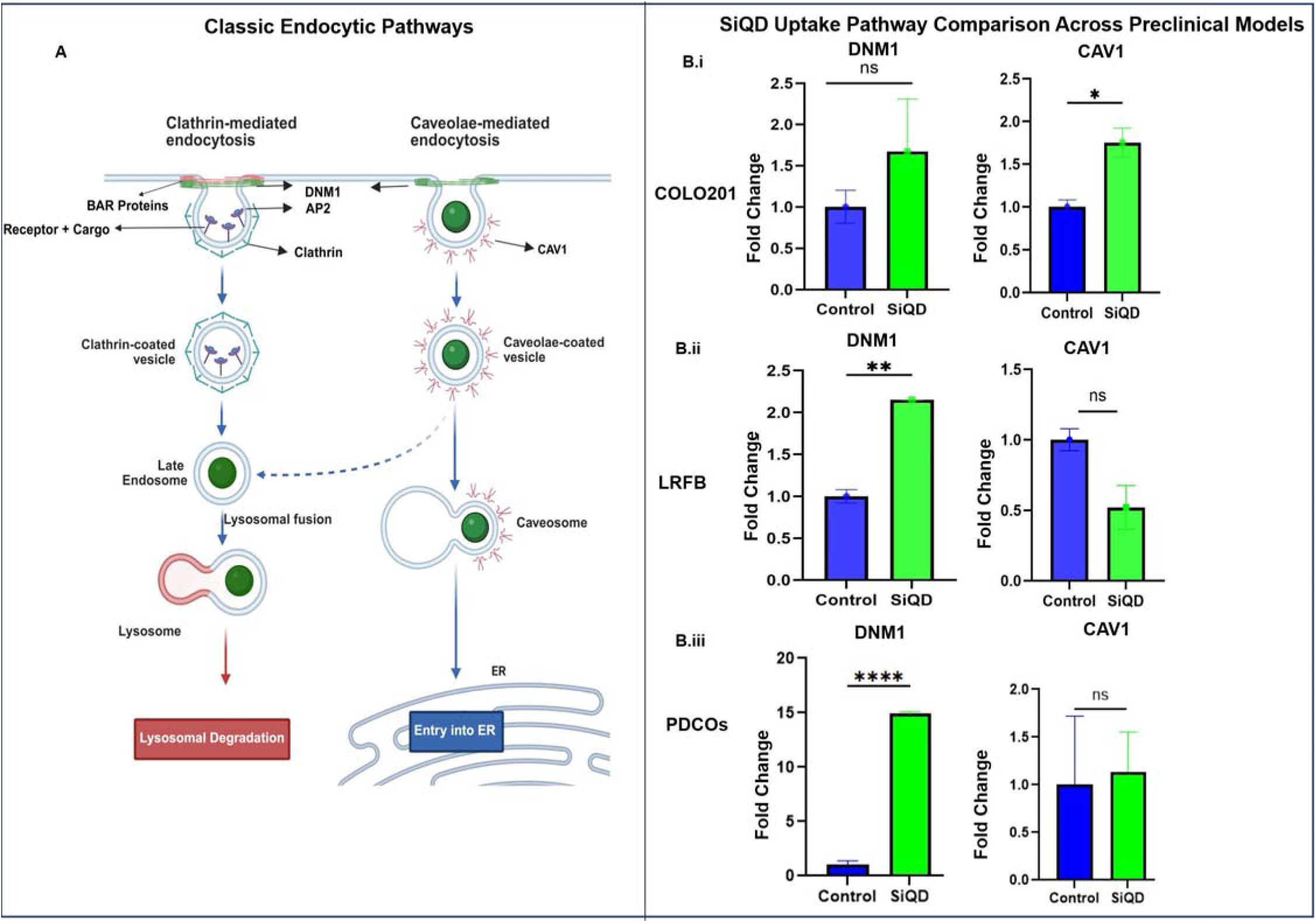
**A** schematic diagram represents classic endocytic uptake pathways of nanoparticles. **Bi,ii,iii** Bargraph representing the comparison of the fold change of CAV1 and DNM1 in COLO201, LRFB and PDCOs.n=3, t-test. Statistical significance: P < 0.05*, P < 0.01**, P < 0.001***,P < 0.00001****

We hypothesized that nanoparticle uptake involves multiple pathways, as extensively reported in the literature ^54, 55^. Instead, we analyzed CAV1 and DNM1 gene expression levels in SiQD-incubated COLO201, LRFB, and patient-derived organoids. Results showed that in COLO201, caveolin-mediated endocytosis is crucial for SiQD uptake, with CAV1 expression increasing 1.7-fold over controls **(Figure 7B. i)**. Conversely, LRFB and PDCOs exhibited a 1.1-and 14-fold increase in DNM1 expression, respectively, suggesting the clathrin pathway as the primary route for SiQD entry, evidenced by a significant increase in fold change **(Figure 7B. ii,7B. iii)**. Overall, the data indicate that clathrin-dependent endocytosis is overactive in PDCO and LRFB cells, leading to SiQD accumulation in lysosomes, whereas in COLO201 cells, the caveolin pathway is upregulated. This suggests SiQDs may localize to lysosomes or endosomes, warranting further study. Additional uptake pathways may also deserve further research.

Our data demonstrates that observed patterns are consistent with involvement of classic nanoparticle uptake pathways. Overall, this research indicates that the intracellular fate of SiQDs varies by cell type. Consequently, this variability can greatly impact nanoparticle applications. Such differences hinder predictable nanoparticle performance and pose a major obstacle for clinical translation. Hence, the study highlights the importance of employing physiologically relevant models to better understand nanoparticle behavior in patients.

This study is a proof of concept and uses only one colorectal cancer cell line and a patient-derived organoid model. This study has shown a significant difference in nanoparticle interaction between cell lines and PDCOs. Therefore, future studies are needed to validate these findings across a large set of patient-derived organoids and colorectal cancer subtypes. This study only focused on comparing two common pathways. Further investigations into pathways such as fast endophilin-mediated endocytosis (FEME) and macropinocytosis would improve understanding of SiQD internalization. This study identified only two TME-related proteins, WNT and EGF, and focused exclusively on the CD44v9-positive chemoresistant cell population. Future studies will investigate a broader range of proteins and evaluate their interactions with different subpopulations of cells in the tumor microenvironment to develop targeted nanoparticle delivery tailored to different cell populations in colorectal cancer.

## Conclusion

In conclusion, this study addresses why nanoparticles that succeed in conventional cell culture so often fail to translate in patients, tracing this gap to the biological identity that nanoparticles acquire through protein corona formation in the tumor microenvironment (TME). Using graphene and silicon quantum dots as material-distinct probes, we demonstrate that nanoparticle identity, uptake, and intracellular fate are mutually governed by material composition and the corresponding biological environment. By profiling nano-bio interactions across fibroblasts, cancer cells, and patient-derived organoids (PDCOs), we show that traditional two-dimensional cell culture overestimates internalization relative to physiologically relevant ex vivo models, establishing the need to incorporate PDCOs in preclinical screening to accurately predict and optimize nano-bio interactions. Critically, TME-specific proteins in the organoid environment drive the selective accumulation of nanoparticles within distinct organoid subpopulations, and engineering the protein corona with the TME-related proteins WNT and EGF redirected SiQD to CD44v9-positive chemoresistant cells, increasing targeted uptake from 3.8% to 70%. We further show that the internalization pathway is model-dependent, PDCOs and fibroblasts favor clathrin/dynamin-mediated endocytosis, whereas cancer cells engage caveolae-mediated entry, influencing whether a nanocarrier is routed toward lysosomal degradation or delivered intact.

Overall, this study establishes PDCOs as a physiologically relevant model for understanding tumor microenvironment-driven nano-bio interactions and provides a framework for manipulating the biological identity of nanoparticles through a TME protein-based protein corona to control their uptake, targeting, and intracellular fate. By transforming the protein corona from an obstacle into a programmable design element, this work positions PDCOs as a next-generation platform for precision nano-oncology and accelerates the development of clinically translatable nanomedicines for targeted drug delivery, molecular imaging, and precision therapeutics.

## Methodology

### Nanoparticle Preparation

1mg of SiQD (ACS material, BASIB001 ) is added to 1 ml Milli-Q water. The solution was then vortexed for 16 hr to completely dissolve the SiQD. GQD (ACS material, GNQD0201), was already suspended in water. TEM images from the manufacturer confirm the size to be less than < 20nm.

### Protein Corona isolation

1 ml of SiQD and GQD were incubated in cancer cell media for 90 mins. To remove the soft protein corona, the incubated samples were centrifuged at 12000 RPM for 45 minutes. The supernatant was removed, and the samples were resuspended in 100 µL Dulbecco’s Phosphate - buffered saline solution (DPBS) (Corning, 21-031-CV).

### Transmission Electron Micrograph

The isolated protein corona was further diluted 1:50 in DPBS. To analyze the protein corona, a FEI Tecnai G2 Spirit BT Transmission Electron Microscope of 100kv voltage and 43000x magnification was performed by the Office of Research and Partnership’s ORP Imaging Cores – Electron Facility at the University of Arizona.

### Time-Dependent Analysis

GQD (ACS material, GNQD0201), and SiQD (ACS Material, BASIB001) samples were incubated in Fetal Bovine Serum (FBS)(ATCC) (1:1) dilution at each time point. The time points are at 15-minute intervals up to 120 minutes. At each point, the protein corona is isolated at 15000 RPM for 45 minutes, resuspended in DPBS, and analyzed for protein concentration using a Nanodrop One^c^ (Thermo Scientific) at A260.

### Size Dependent Analysis

SiQD and GQD protein corona isolated from organoid media and cancer cell media was isolated and further diluted in DPBS 1:1 dilution, and dynamic light scattering was performed in Malvern Zeta sizer DLS machine, n= 3.

### Fourier Transform - Infrared Spectroscopy

The isolated SiQD and GQD protein coronas were spread on a silicon wafer. Thermo Fisher Scientific Nicolet IS50R FT-IR with a liquid-nitrogen-cooled, high-sensitivity MCT-A detector (11,700–600 cm-1) was performed on the sample. Origin Pro and Python were used to perform further analysis.

### Generation and Culture of Human Patient-Derived Colonic Organoids (PDCO) Two-Dimensional Monolayer System from Tumor Tissues

Human colorectal cancer 2D monolayer system developed from patient tissues were generated based on the previously published article^37^. Tissues were minced using surgical scalpel blades on a cell culture Petri dish. The fragmented tissues were washed with ice-cold DPBS + Pencillin/ Streptomycin (Fisher Scientific). Small chunks of tissue were transferred into basal media supplemented with 10 μM Y-27632 (Stem Cell Technologies) and 1mg/mL Collagenase Type 3 (MP Biomedicals) for digestion. Tissues were incubated at 37 °C for 45-60 minutes in an orbital shaker. Further, TrypLE(Fisher Scientific) + 10 μM Y-27632 were added to the undigested tissue and incubated at 37°C for 45-60 minutes. The digested clusters were combined and centrifuged at 500 × g for 5 minutes at 4°C to pellet cells. The supernatant was discarded, then the pellet was resuspended in ice-cold DPBS + AB. The sample was centrifuged at 500 × g for 5 minutes at 4 °C. The supernatant was discarded, and the cell pellets were resuspended in an appropriate volume of ice-cold Matrigel®(Corning). Further 25 µl cell-Matrigel® droplets were seeded in prewarmed 48-well tissue culture plates. Incubate the cell-Matrigel® droplets at 37 °C for 20 minutes to solidify as a dome, then overlay with growth media as mentioned in **Supplementary Table 1**. Incubate the 3D organoids until they are fully mature.

Organoid media was removed from 3D organoids, and using ice-cold advanced DMEM/F12, 3D organoids were harvested and centrifuged at 500 × g for 5 min. After centrifugation, pellets were collected and resuspended in advanced DMEM /F12, and the above steps were performed 3 – 4 times until Matrigel was removed. Pellets were further resuspended in 0.25% trypsin-Y27 and incubated at 37°C for 3-5 minutes. 100% FBS was added to quench the trypsin activity. The sample was centrifuged at 4 °C, 500 × g for 10 minutes. Supernatants were removed, and pellets were resuspended in seeding media (Supplementary Table 1). 10^5^ cells were seeded into the Matrigel-coated plate. Plates were further centrifuged at 300 × g for 1 minute at 37°C. Cells were cultured in seeding media at 37°C, 5% CO_2_ for 16-24 hours. After 16-24 hours, gently wash wells with advanced DMEM/F12 and add complete LWRN to the well. Incubate at 37°C 5% CO_2_.

### Cell Culture

Human cell lines: fibroblast (LRFB) and colon cancer cell line (COLO201, (ATCC) were trypsinized for 3mins and further centrifuged at 250 RPM for 12min and further cultured in Advanced Dulbecco Modified Eagle Medium (DMEM) / F12 (Gibco) and RPMI-1640 (CORNING), respectively, with 10% FBS (ATCC) and 5% Antibiotic Antimycotic Solution (CORNING).

### Fibroblast Immortalization

Fibroblasts were disassociated from a biopsy of normal colonic tissue. Tissue was minced in small pieces, digested with Collagenase 3 for 15 min, followed by 15 min with 0.25% Trypsin. Once dissociated, cells were plated onto a 24 well and grown in DMEM with 20% FBS to select for fibroblasts and remove epithelial cells. After 2 days of growth, fibroblasts were immortalized by overexpressing hTERT and CDK4 via lentiviral infection. Prior to infection, virus had been concentrated to 4x with LentiX Concentrator (Takara Bio, 631232). 250uL of each virus (hTERT and CDK4 made separately) was diluted into 2 mL of DMEM + 20% FBS media and added onto the fibroblasts for 48 hours. Once confluent, the fibroblasts were removed from the 24 well plate into a 10cm plate where they were maintained in DMEM + 20% FBS + 1% P/S.

### Spectral Flowcytometry

After incubating SiQD and GQD with 10^5 cells LRFB, COLO201 and PDCOs, the samples will be trypsinized using trypsin (CORNING) for 3 minutes, then Advanced DMEM/F12 (Gibco) with 10% FBS (ATCC) will be added to halt the reaction. Next, the samples will be centrifuged at 250 RPM for 12 minutes at 4°C. The supernatant will be discarded, and the cell pellet will be resuspended in DPBS(CORNING).BD FACS CANTO was used to analyze the sample. Further analysis was performed using Omiq^56^. Gating strategies **(Supplementary Figure 13)** were applied, and graphs were plotted using GraphPad Prism (GraphPad Software, San Diego, CA).

### Immunofluorescence

SiQD (1mg/ml) was incubated in organoids for 16 hours, and additional organoids were fixed with 4% paraformaldehyde (PFA) at 37°C. After fixation, PFA (Thermo Fisher Scientific, 28908) was removed, and the organoids were washed three times with 1X PBS. 0.2% Triton X-100 (Thermo Fisher Scientific, A16046.AE) was used at RT to permeabilize the cells. After permeabilization, organoids were washed three times with PBS, and 2.5% BSA (Millipore Sigma, A7030) in PBS was added as the blocking buffer. After blocking, CD44v9 (1:1000) (Cosmo Bio, CAC-LKG-M001) was added and incubated for 2 hr at RT in the dark. Later, PBST was used for washing, and Anti-Rat (Thermo Fisher Scientific, A21247) was added as a secondary antibody. Further, 2 µM DAPI (Thermo Fisher Scientific, D1306) and Phalloidin (Thermo Fisher Scientific, A22283) were mixed in PBS and incubated for 30 min. Afterward, 100 µL of PBS was added, and the organoids were imaged using a Nikon SoRa Spinning-Disk Confocal Microscope. Further, images were processed using Nikon Elements.

### Polymerase Chain Reaction

Organoid, LRFB, and COLO201 cells were treated with SiQD. RNA was extracted with the PureLink™ RNA Mini Kit (Invitrogen). The transcripts of Caveolin 1 (CAV1), Dynamin 1 (DNM1), and Actin B (ACT B) (Housekeeping gene) were amplified using the TaqPath™ 1-Step Multiplex assay. Initially to prepare the master mix for the gene expression analysis, TaqPath™ 1-Step Multiplex Master Mix (4X) (applied biosystems), primers (CAV1 FAM (ThermoFisher Scientific Hs00971716_m1), DNM1 FAM (ThermoFisher Scientific, Hs01074761_m1) and ACT B FAM (ThermoFisher Scientific, Hs01060665_g1) 1 µL/assay Use primer concentrations of 150–900 nM and a probe concentration of 100–250 nM. RNA Sample (100ng) and RT-PCR-grade water were used to fill the total reaction volume of 20 µL. Applied Biosystems™ real-time PCR instrument was used to run the analysis with the following thermal cycle conditions. Analysis was performed by normalizing the samples to ACTB gene expression. Replicated details were provided in the figure panel.

| Step | Stage | Cycles | Temperature | Time |
| --- | --- | --- | --- | --- |
| UNG incubation | 1 | 1 | 25°C | 2min |
| Reverse transcription | 2 | 1 | 53°C | 10min |
| Polymerase activation | 3 | 1 | 95°C | 2min |
| Amplification | 4 | 40 | 95°C | 3s |
|  |  |  | 60°C | 1min |

### Data Analysis

Significance of the results was tested using commercially available software (GraphPad Prism, GraphPad Software, San Diego, CA). Analysis was performed using a t-test, where a p-value of less than 0.05 was determined significant. Details of biological replicates are mentioned in figure captions.

## Author Information

### Corresponding Authors

Swarna Ganesh - Assistant Professor, Department of Biomedical Engineering, University of Arizona College of Medicine, Tucson, AZ, email ID

### Author Contributions

M.P.R. and S.G. conceived and designed the study. M.P.R. conducted the experiments, performed the data analysis, and prepared the original manuscript draft. K.W.P. performed image acquisition using the Nikon SoRa spinning-disk confocal microscope. J.C. established the patient-derived organoid two-dimensional monolayer system. S.G., K.W.P., and J.C. reviewed the manuscript.

## Acknowledgements

We thank the members of the Ganesh Lab for their valuable discussions and helpful feedback throughout this work. This work was supported by the University of Arizona Cancer Engineering Initiative, and in part by the State of Arizona Technology Research and Innovation Fund (TRIF) and the BIO5 Institute

Tissue acquisition was supported by the Tissue Acquisition and Cellular / Molecular Analysis (TACMASR) Core at the University of Arizona, grant: NIH CA023074. Biology Development and Research Organoids (BIODROid) developed patient-derived organoids with support from the cancer center grant (P30 CA023074). Fibroblast cell line was immortalized and provided by Lauren Riede. Patient-derived organoids and organoid growth media resources were generously provided by the Pond Lab. Imaging was performed in part through the Nikon Center of Excellence, Microscopy Shared Resource at the University of Arizona Cancer Center.

Flow Cytometry was supported by the University of Arizona Flow Cytometry and Immune Monitoring Shared Resource, funded by the National Cancer Institute of the National Institutes of Health under award number P30 CA023074.RRID: SCR_023432

We thank Paula Tonino, Manager of the Office of Research and Partnerships Imaging Cores – Electron Microscopy Facility at the University of Arizona (RRID: SCR_023279), for providing transmission electron microscopy (TEM) imaging services.

Fourier transform infrared (FTIR) spectroscopy was supported by the University of Arizona CBC W. M. Keck Center for Nano-Scale Imaging (RRID: SCR_022884).

