## Supplementary Information for "Patient-Derived Organoids as a Model to Understand Tumor Microenvironment-Driven Nano-Bio Interactions – A Framework Towards Improved Nanomedicine Translation"

**Corresponding author (**
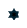
**):**

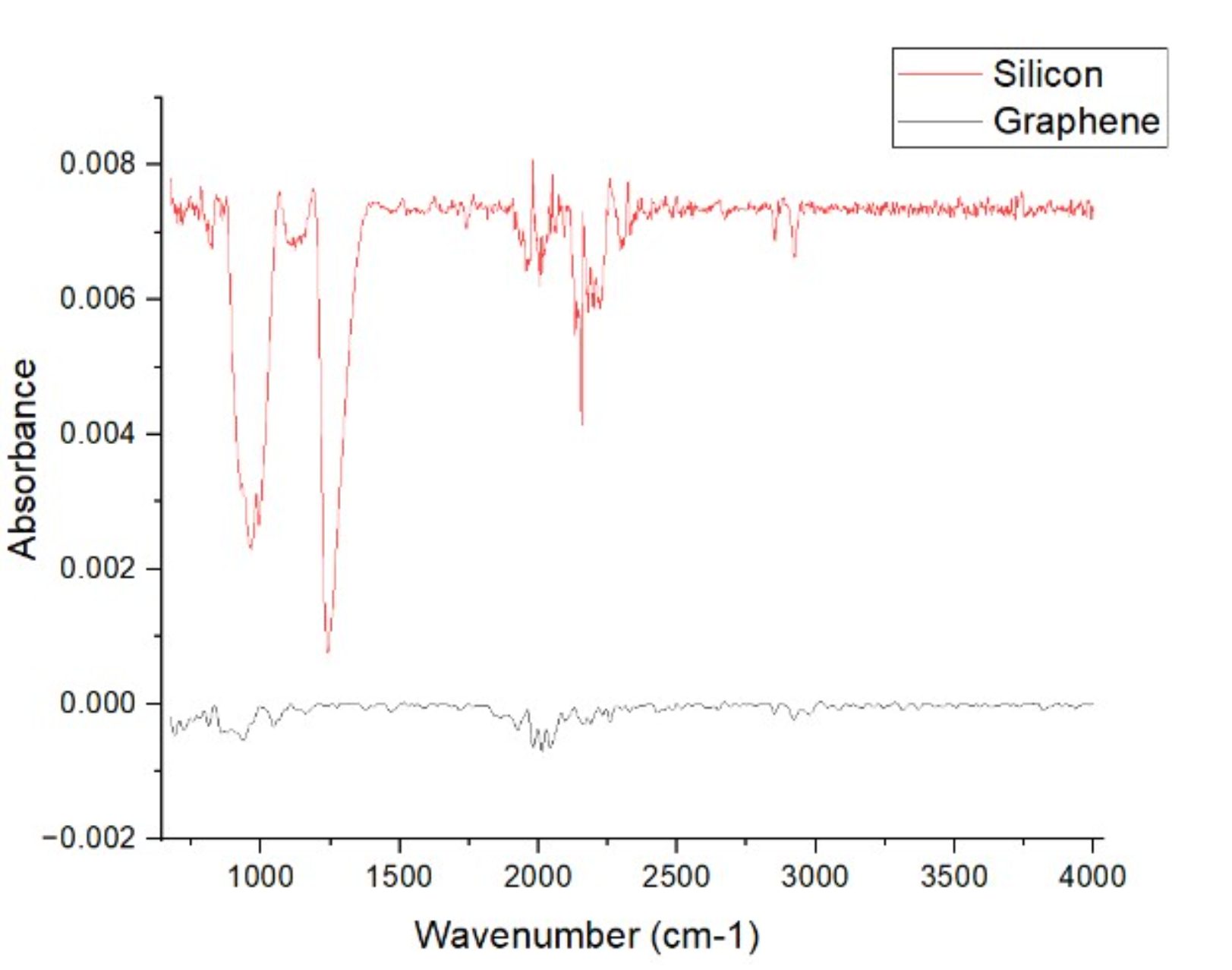
 **Supplementary Figure 1:** FT-IR spectra representing the surface chemistry of the SiQD and GQD.

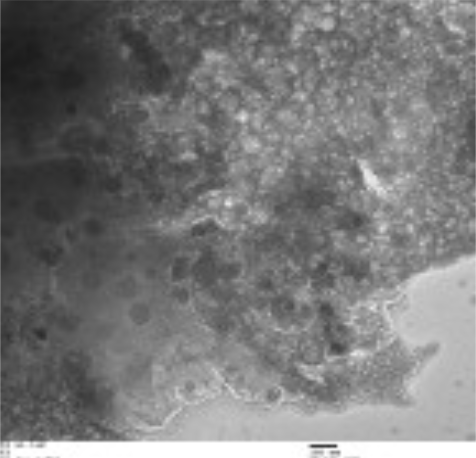

**GQD Protein corona**

**SiQD Protein corona**

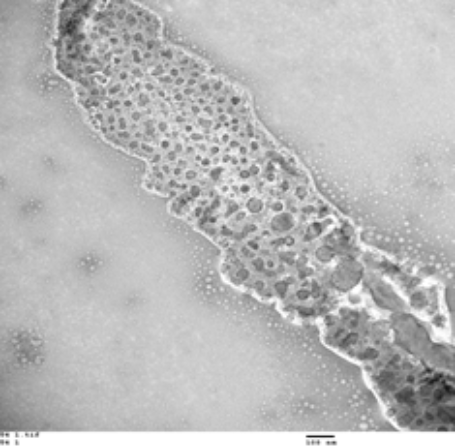

**Supplementary figure 2:** TEM images of SiQD ( X =200nm), GQD(X=20nm) and SiQD protein corona (X=100nm) and GQD protein corona (X= 100nm).

| **Reagent** | **Manufacturer / Catalog Number** |
| --- | --- |
| LWRN conditioned media | In-house |
| Advanced DMEM/F12 | Thermo Fisher — 12634010 |
| HEPES | Corning — 25-060-CI |
| GlutaMAX | Gibco — 35050-061 |
| NAC (N-Acetyl Cysteine) | SIGMA — A9165 |
| Primocin | InvivoGen — ant-pm-1 |
| rhEGF | Invitrogen — BMS320 |
| SB202190 | StemCell — 72632 |
| A83-01 | Tocris — 2939 |
| N2 MAX supplement | R&D — AR009 |
| N21 MAX supplement | R&D — AR008 |
| Nicotinamide | SIGMA — N3376 |
| Pen/Strep | Corning — 30-002-CI |

**Supplementary Table 1:** Colorectal cancer patient-derived organoid media components.

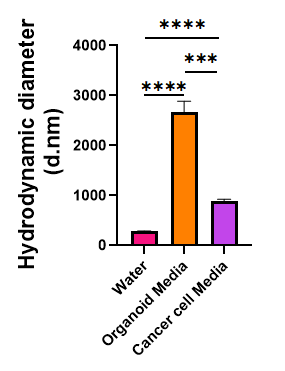

**Hydrodynamic Diameter (nm)**

**Supplementary Figure 3:** The bar graph represents the hydrodynamic diameter of the SiQD in water, organoid media, and cancer cell media, t test,n=3, Statistical significance: P < 0.05
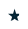
 , P < 0.01
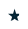

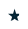
, P < 0.001
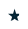

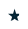

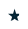
,P < 0.00001
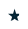

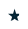

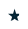

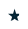

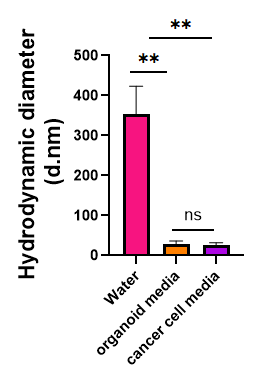

**Hydrodynamic Diameter (nm)**

**Supplementary Figure 4:** The bar graph represents the hydrodynamic diameter of the GQD in water, organoid media, and cancer cell media, t test,n=3, Statistical significance: P < 0.05
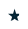
 , P < 0.01
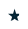

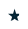
, P < 0.001
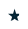

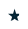

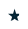
,

P < 0.00001
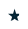

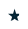

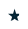

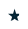

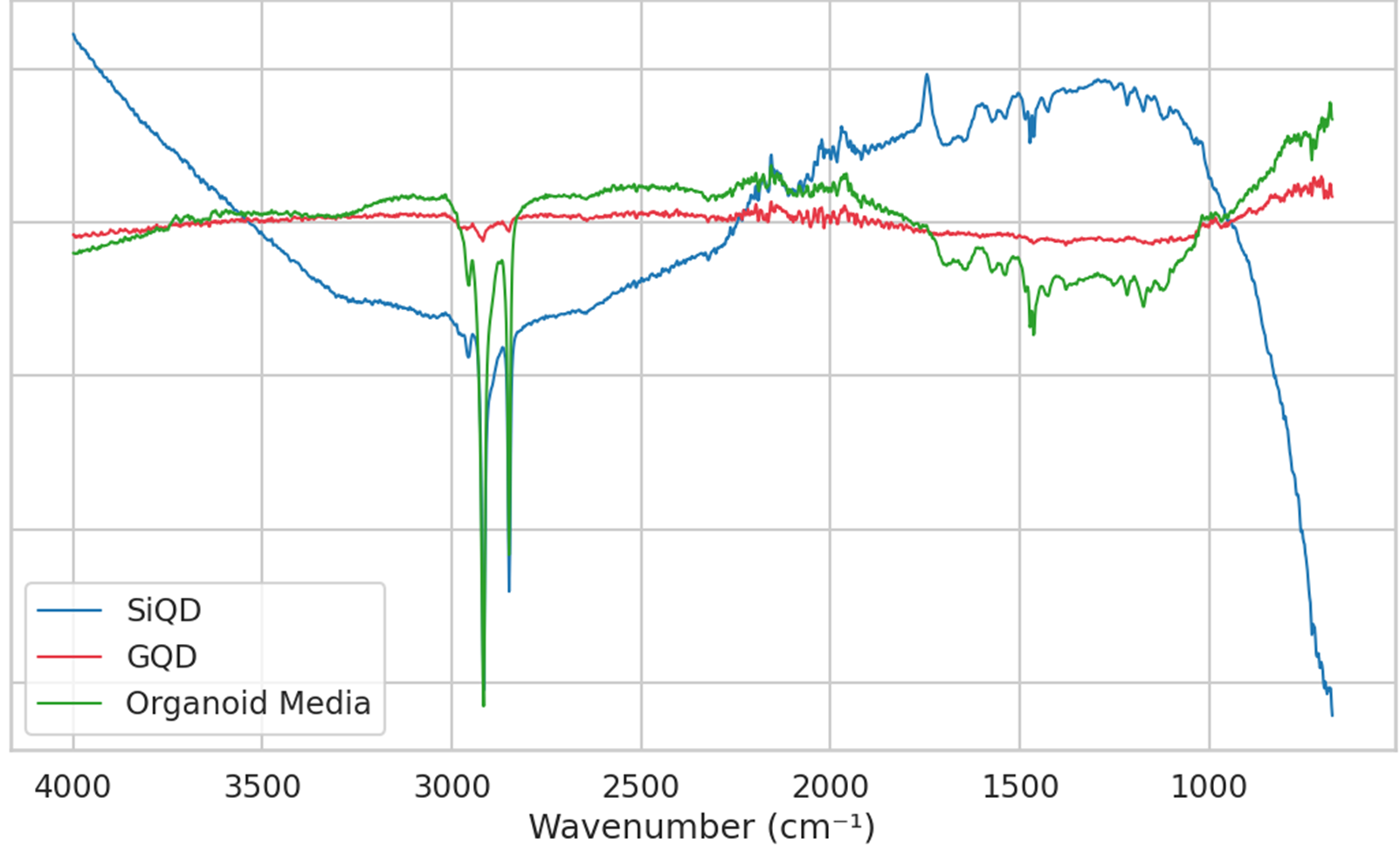

**Supplementary Figure 5:** FT- IR spectra shows the surface chemistry of the protein corona formed by SiQD and GQD in organoid media and organoid media as a control, n=3.

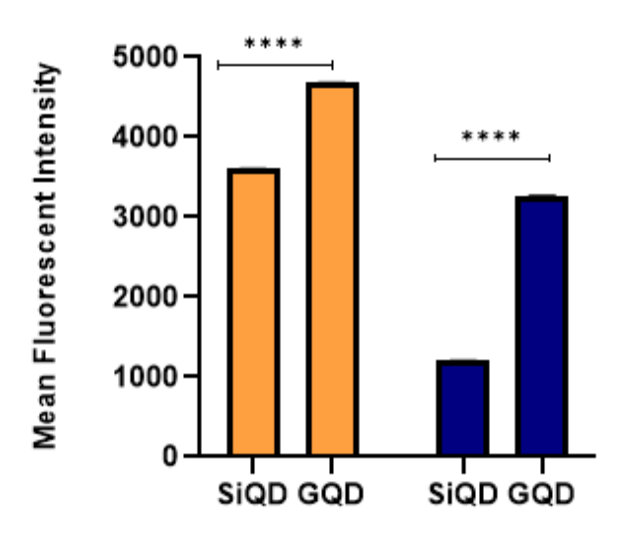

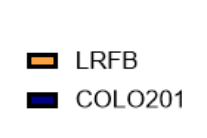

**Supplementary Figure 6:** Barplot representing the mean fluorescent intensity of the SiQD and GQD in LRFB and COLO201, n= 10,000 cells , t test. , Statistical significance: P < 0.05
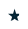
 , P < 0.01
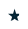

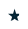
, P < 0.001
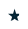

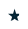

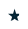
,

P < 0.00001
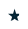

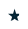

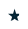

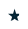

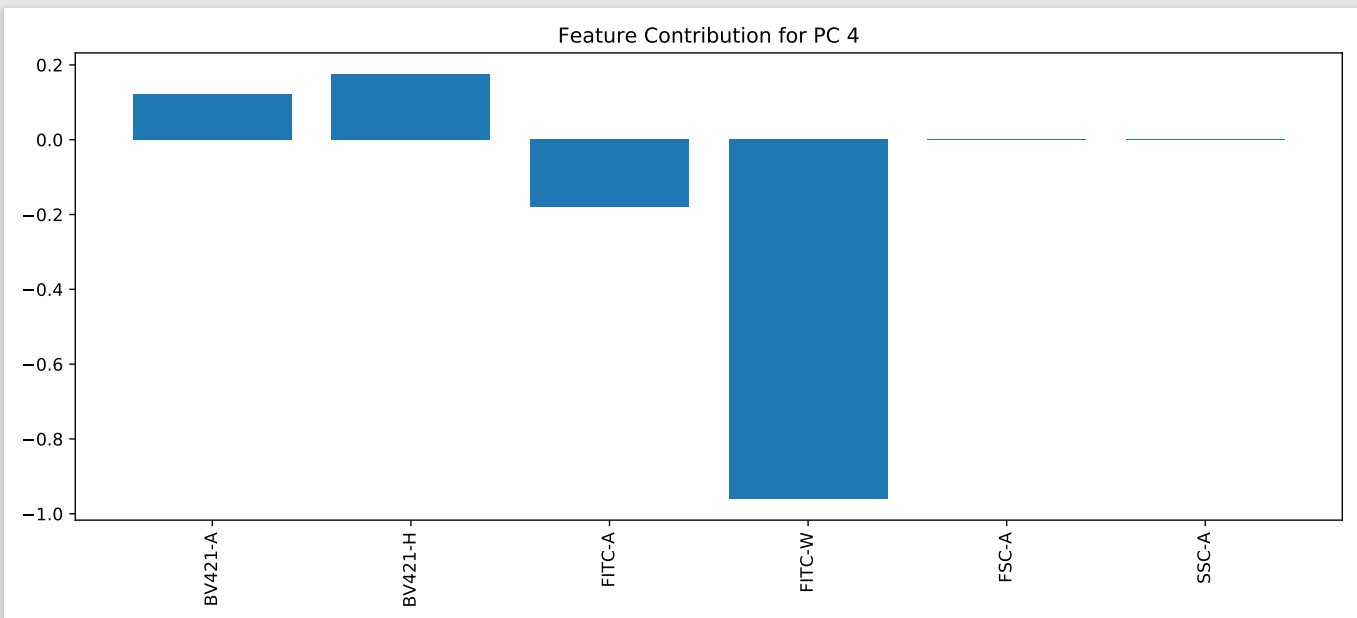

**Supplementary Figure 7:** Feature Contributions to principal component 4 by fluorescence of SiQD and GQD in LRFB, n=10,000 cells.

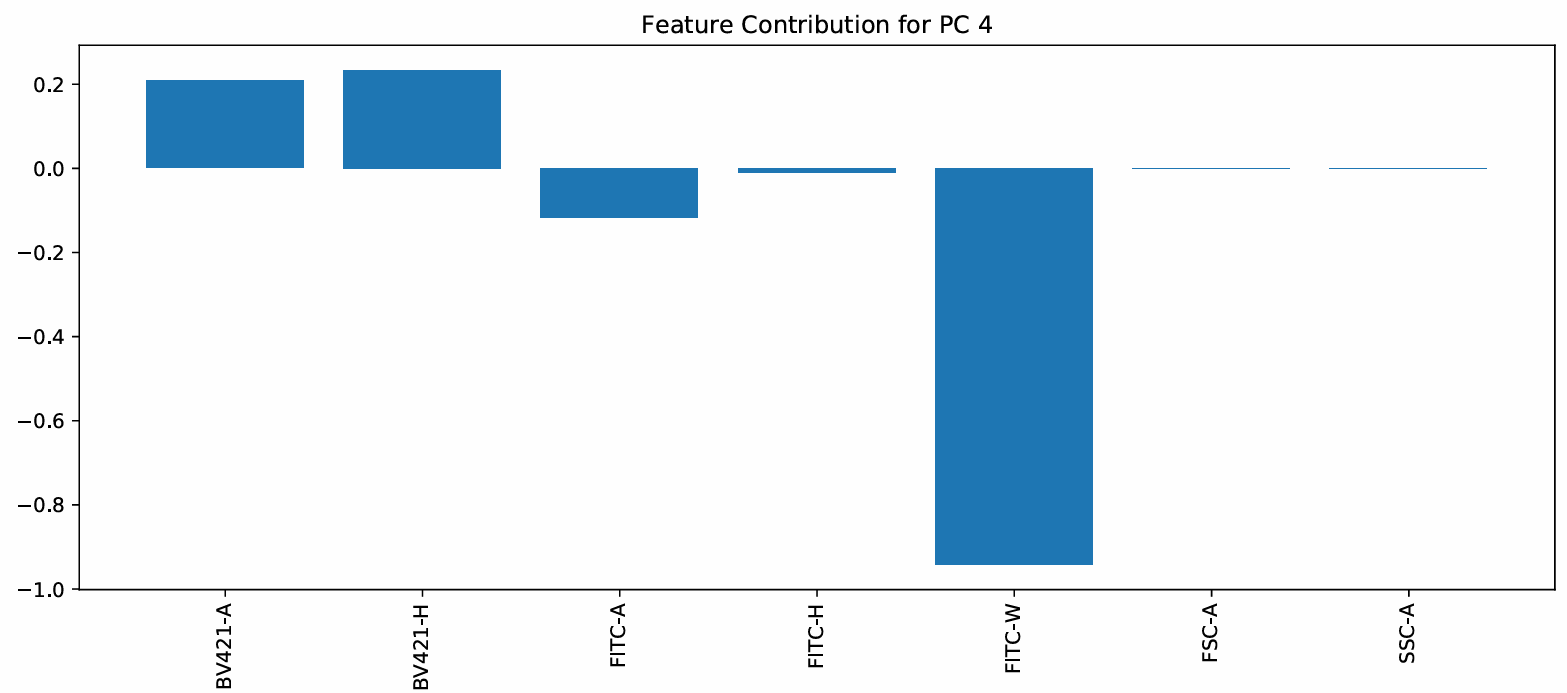

**Supplementary Figure 8:** Feature Contributions to principal component 4 by fluorescence of SiQD and GQD in COLO201, n=10,000 cells.

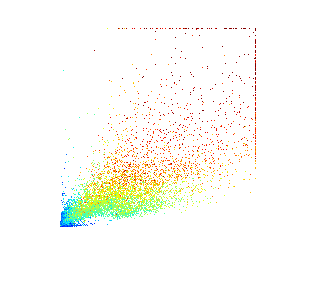

FSC-A

SSC-A

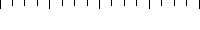

0.0

66K

131K

197K

262K

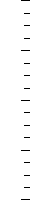

0.0

66K

131K

197K

262K

FITC-A

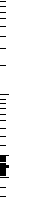

0

10²

-10²

10³

**Supplementary Figure 9**: Scatter plot representing the SiQD uptake in PDOs**,** n= 10,000 cells.

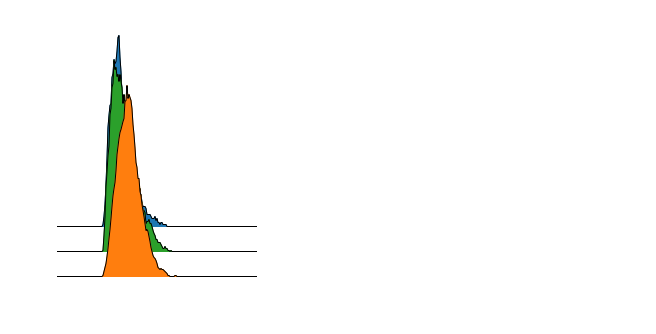

FITC-A

count

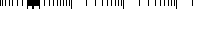

0

10³

10⁴

10⁵

6hr

48hr

24hr

**Supplementary Figure 10:** Histogram comparing the mean fluorescent intensity of SiQD uptake in PDOs at 6 hr, 24 hr, and 48 hr, n= 10,000 cells

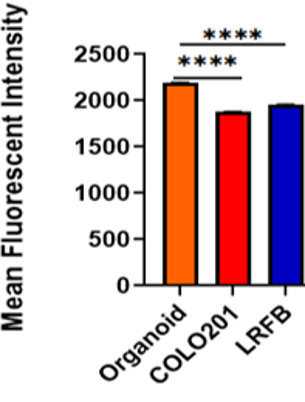

**Supplementary Figure 11**.Barplot compares mean fluorescent intensity across LRFB, COLO201 and PDOs, n= 10,000 cells,t test

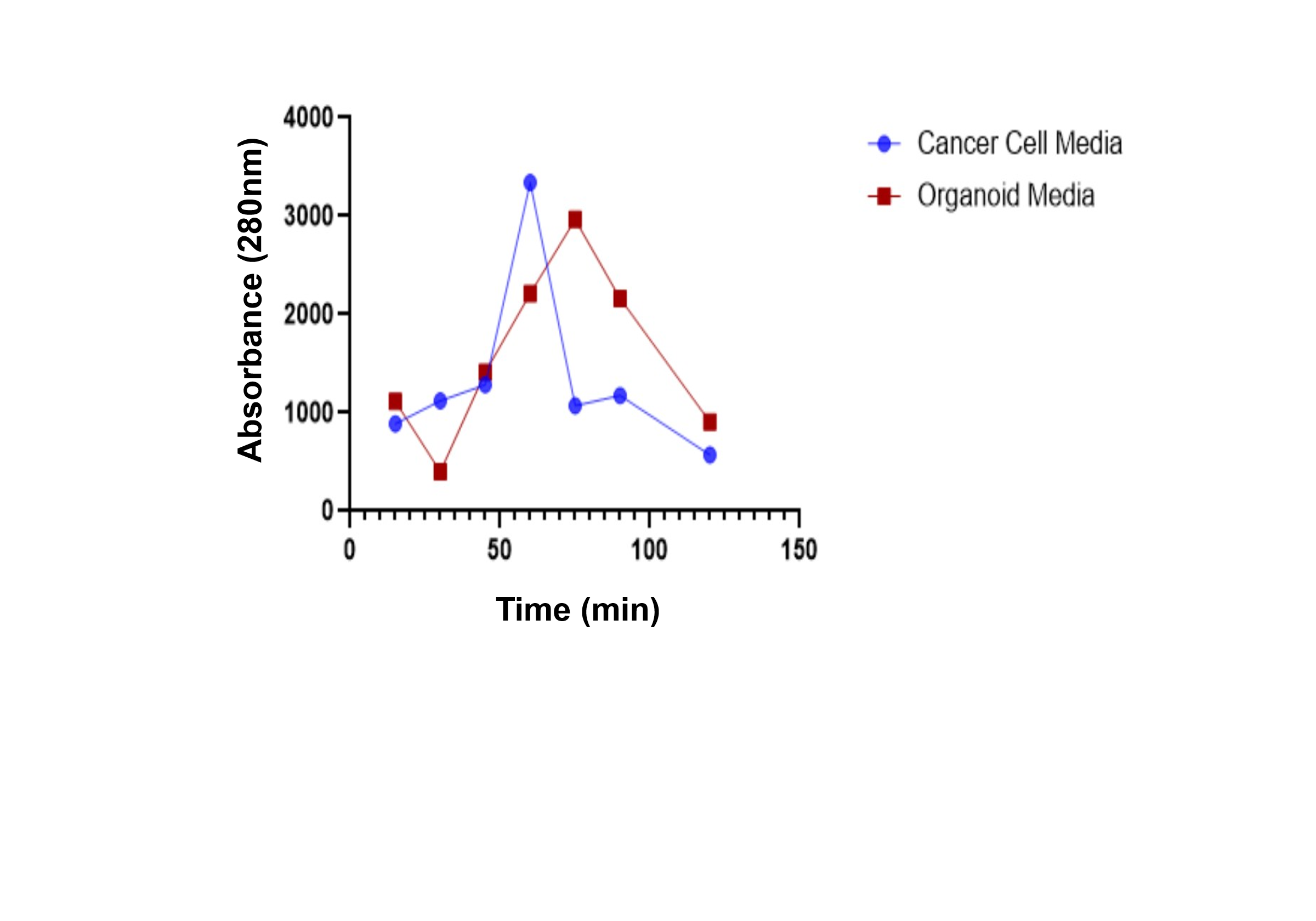

**Supplementary Figure 12:** Protein corona formation by SiQD in Cancer cell media and organoid media, X-axis = Time (min), Y-axis = Absorbance at 280 nm.

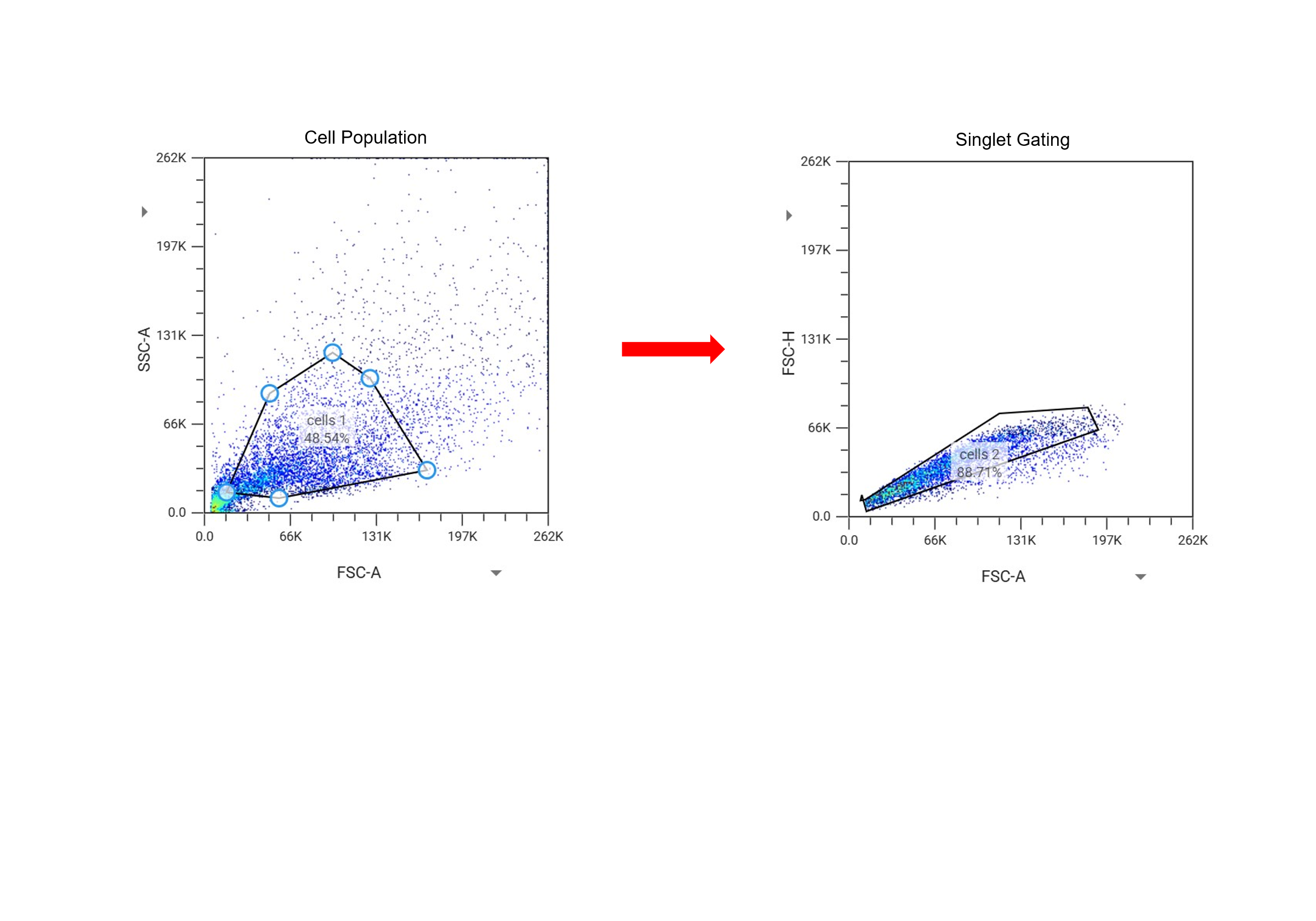

**Supplementary Figure 13:** Gating strategies for PDO and cell uptake spectral flow cytometry data

.
